# Cleavage of cFLIP restrains cell death during viral infection and tissue injury and favors tissue repair

**DOI:** 10.1101/2022.12.15.520548

**Authors:** Kristel Martinez Lagunas, Deniz Pinar Savcigil, Matea Zrilic, Carlos Carvajal Fraile, Andrew Craxton, Emily Self, Iratxe Uranga-Murillo, Diego de Miguel, Maykel Arias, Sebastian Willenborg, Michael Piekarek, Marie Christine Albert, Kalvin Nugraha, Ina Lisewski, Erika Janakova, Natalia Igual, Wulf Tonnus, Ximena Hildebrandt, Mohammed Ibrahim, Marlies Ballegeer, Xavier Saelens, Andrew Kueh, Pascal Meier, Andreas Linkermann, Julian Pardo, Sabine Eming, Henning Walczak, Marion MacFarlane, Nieves Peltzer, Alessandro Annibaldi

**Author notes:** These authors contributed equally to this work (Kristel Martinez Lagunas, Deniz Pinar Savcigil).

## Abstract

Cell death coordinates repair programs following pathogen attack and tissue injury. However, aberrant cell death can interfere with such programs and cause organ failure. cFLIP is a crucial regulator of cell death and a substrate of Caspase-8. Yet, the physiological role of cFLIP cleavage by Caspase-8 remains elusive. Here, we discovered an essential role for cFLIP cleavage in restraining cell death in different pathophysiological scenarios. Mice expressing a cleavage-resistant cFLIP mutant, *Cflip*^*D377A*^, exhibited increased sensitivity to SARS-CoV-induced lethality, impaired skin wound healing and increased tissue damage caused by *Sharpin* deficiency. *In vitro*, abrogation of cFLIP cleavage sensitizes cells to TNF-induced necroptosis and apoptosis by favoring complex-II formation. Mechanistically, the cell death-sensitizing effect of the D377A mutation depends on Gln(Q)469. These results reveal a crucial role for cFLIP cleavage in controlling the amplitude of cell death responses occurring upon tissue stress, to ensure the execution of repair programs.

## Introduction

Cell death is a fundamental biological process that ensures tissue homeostasis and orchestrates tissue remodelling following injury or infection. However, if on the one hand abrogation of cell death responses can prevent the activation of repair programs, on the other hand exacerbated cell death can lead to tissue failure (*1–3*). Therefore, the ability of tissues to control the extent of cell death under stress conditions is of fundamental importance for the activation of optimal repair programs. TNF is a pro-inflammatory cytokine that is produced in response to a large variety of stressors, including viral infection and injury, and it can initiate repair processes by inducing the expression of pro-inflammatory genes or by triggering cell death (*1, 4, 5*). The mechanisms regulating the decision between these two outcomes are of fundamental importance for the maintenance of tissue homeostasis and for the capacity of tissues to overcome damage (*6, 7*). Binding of TNF to TNFR1 results in the formation of two spatially and temporally distinct complexes (*8*). A membrane bound complex, called complex-I, is assembled on the intracellular, death domain (DD) containing portion of TNFR1. It is composed of adaptor proteins, such as TRADD and TRAF2, kinases, such as RIPK1, IKKα and IKKβ, TAK1, TBK1 and IKKε, and E3 ligases, such as cIAP1/2 and LUBAC (*9–14*). The concerted action of phosphorylation and ubiquitination events ensures the correct assembly and stability of the complex, leading to the activation of genes required to mount an inflammatory response (*1, 15, 16*). Any perturbation of phosphorylation or ubiquitination processes leads to the formation of a secondary, cytoplasmic complex, referred to as complex-II (*1, 14, 16–21*). This complex consists of FADD, RIPK1, cFLIP, Caspase-8 and, depending on the cell types, RIPK3 (*18, 22*). Complex-II has cytotoxic activity and can trigger Caspase-8-mediated apoptosis and RIPK1/RIPK3/MLKL-mediated necroptosis (*23*). In immune cells, Caspase-8 activation can result in Gasdermin D cleavage with the consequent induction of pyroptosis (*24*). Apart from TNFR1, other immune receptors including TLR3, TLR4 and type I and type II interferon receptors have the potential to induce a complex-II-like cell death-inducing platform (*25–27*). The cFLIP_L_/Caspase-8 heterodimer acts as a molecular switch that controls the cell death outcomes of complex-II. Indeed, while the cFLIP/Caspase-8 heterodimer is required to suppress necroptosis, the Caspase-8/Caspase-8 homodimer formation is required for the activation of downstream Caspase-3 and −7 and induction of apoptosis (*28–30*). Therefore, cFLIP_L_ (herein referred to as cFLIP) represents both an activator and an inhibitor of Caspase-8, since it is needed for the ability of the latter to suppress necroptosis, but at the same time it prevents Caspase-8 full activation and apoptosis (*28*). cFLIP is a catalytically inactive homolog of Caspase-8, composed of tandem death effector domains (DED) and a caspase-like domain formed of a large and a small subunit (*31*). Apart from being the most direct, non-redundant regulator of Caspase-8 activity, cFLIP is also a Caspase-8 substrate. Caspase-8 can cleave cFLIP at aspartic acid (D) 376 (377 in mouse) (*32*). Different studies have addressed the function of the cleavage at D376 mainly using cell-free systems (*31, 33, 34*), however the biological role of this proteolytic event on cFLIP remains to be elucidated.

Here, we characterized the dynamics and function of cFLIP cleavage at D377 both *in vivo* and *in vitro* by generating a mutant mouse bearing a point mutation that abrogates the cleavage at Asp377 (*Cflip^D377A^* mice). The D377A mutation renders mice more sensitive to the lethal effect of SARS-CoV infection, exacerbates the phenotype of the *Sharpin*^*cpdm*^ mice and impairs skin wound healing. Cells derived from the *Cflip*^*D377A*^ mice are significantly more sensitive to TNF-induced necroptosis and apoptosis and exhibit increased complex-II formation. At the mechanistic level, we observed that Glutamine Q469, a residue important for cFLIP heterodimerization with Caspase-8, is responsible for the cell death sensitizing effect of the cFLIP_D377A mutant. Therefore, we report on the precise biological function of cFLIP cleavage in keeping cell death responses in check during pathogen attack, tissue injury and damage, to favor remodelling and repair.

## Results

### cFLIP cleavage by Caspase-8 limits TNF cytotoxicity

We first characterized the dynamics of cFLIP cleavage in TNF signalling. TNF treatment alone (T) or in combination with Smac mimetic (S) and/or emricasan (E), rapidly induced cleavage of cFLIP, as demonstrated by the appearance of the p43 fragment (Fig. 1A). Of note, cFLIP cleavage occurs in the absence of full Caspase-8 activation, as in the case of TNF-treatment alone, and despite the inhibition of Caspase-8 activity (Fig. 1A and 1B). We next addressed the molecular requirements for cFLIP cleavage and we found that FADD and Caspase-8, but not Caspase-3/7, are necessary for cFLIP to be cleaved (Fig. 1C, 1D and S1A, S1B, S1C and S1D). Since the co-deletion of Caspase-3 and 7 did not affect the cleavage event (Fig. 1C), we conclude that upon TNF stimulation, the formation of a FADD/Caspase-8 complex is required for cFLIP cleavage and that Caspase-8 is most likely the only caspase able to cleave cFLIP in mouse cells. cFLIP bears an aspartic acid residue in close proximity to D377, the D371, that could potentially undergo Caspase-8-mediated cleavage (Fig. 1E) (*35*). In order to elucidate whether D371 can substitute for D377A, we reconstituted *Cflip^-/-^* MDFs with either WT cFLIP or mutant cFLIP bearing substitutions in D371A or D377A (Fig. S1E). We found that whereas D371 is dispensable for cFLIP cleavage, the D377 is the only cleavage site of cFLIP upon TNF signalling pathway activation (Fig. 1E). Re-expression of the WT or mutant cFLIP variants could protect the *Cflip^-/-^* MDFs from TNF-induced cell death (Fig. S1F). Intriguingly, the cleavage-deficient D377A mutant cFLIP sensitized MDFs to TNF and emricasan-induced necroptosis (Fig. 1F). These findings indicate that cFLIP is cleaved by Caspase-8 following TNF-signalling pathway stimulation at residue D377 and that this cleavage event has pro-survival functions.

**Fig. 1.**
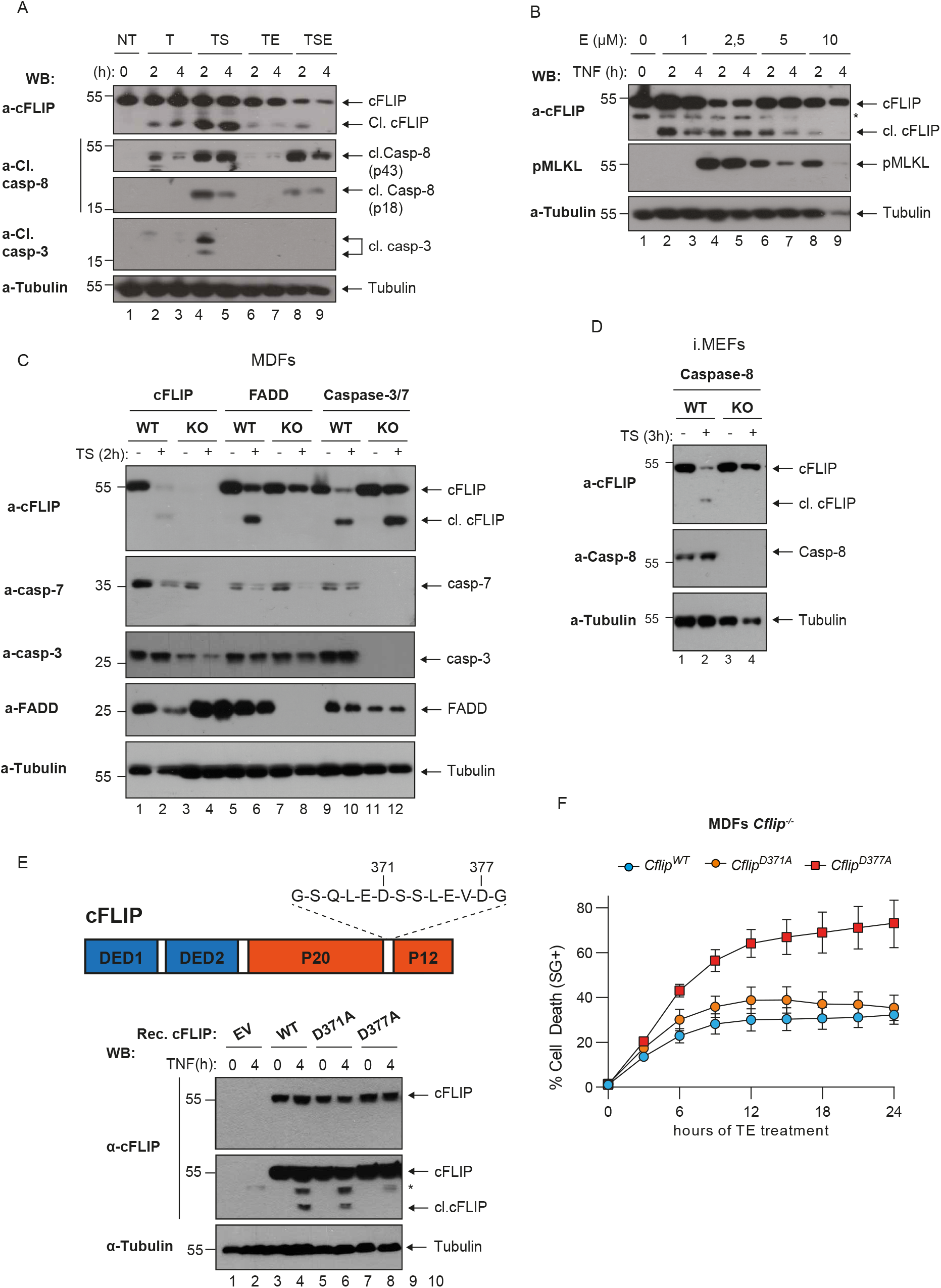
cFLIP is cleaved at Asp377 following TNFR1 activation. (**A**) WT mouse dermal fibroblasts (MDFs) were treated with TNF (T, 100 ng/ml), TNF and Smac mimetic (250 nM, TS), TNF and emricasan (1 μM, TE) and TNF, Smac mimetic and emricasan (TSE) with the indicated time points. Cell lysates were analyzed by immunoblotting for the indicated specific antibodies (n=2). (**B**) WT MDFs were treated with TNF (100 ng/ml) and increasing concentration of emricasan for the indicated time points. Cell lysates were analyzed by immunoblotting with the indicated specific antibodies (n=3). (**C**) MFDs and (**D**) mouse embryonic fibroblasts (MEFs) of the indicated genotypes were treated with TNF (100 ng/ml) and Smac mimetics (250 nM) for two hours and cell lysates analyzed by immunoblotting with the indicated specific antibodies (n=2). (**E**) Cartoon depicting cFLIP domain composition and aspartic acid residues 371 and 377 (upper part). Cflip^-/-^ MDFs were reconstituted either with empty virus or virus expressing WT or the indicated mutant versions of cFLIP and treated with TNF (100 ng/ml) for 4 hours. Cell lysates were analyzed by immunoblotting with the indicated specific antibodies (n=2). (**F**) Cells as in E were treated with TNF (10 ng/ml) and emricasan (1 μM) and cell death was measured over time by calculating the percentage of Sytox Green positive cells (n=4).

### Non-cleavable cFLIP sensitizes to TNF-induced necroptosis and apoptosis

To study the physiological function of cFLIP cleavage at D377, we generated cFLIP cleavage resistant mice, referred to as *Cflip*^*D377A*^ mice, where the aspartate 377 was mutated to alanine (Fig. S2A). The *Cflip*^*D377A*^ mutant mice were weaned at the expected mendelian ratio and do not exhibit any overt phenotype (Fig. S2B, S2C and S2D). These mice had an overtly normal immune system (Fig. S2E). Mouse dermal fibroblast (MDFs) isolated from WT and cFLIP-mutant mice had comparable levels of cFLIP, and in the D377A mutant cells, cFLIP could not undergo proteolytic cleavage following TNFR1 pathway activation (Fig. 2A). Next, we tested the role of cFLIP cleavage in TNF-induced cell death. While *Cflip*^*D377A*^ cells were not sensitive to TNF treatment alone (Fig. S3A), MDFs, lung endothelial cells (LECs), lung fibroblasts (LFs) and bone marrow derived macrophages (BMDMs) were significantly more sensitive to TNF and emricasan (TE)- or TNF, emricasan and birinapant (TBE)-induced necroptosis, compared to their WT counterparts (Fig. 2B, 2C, 2D and 2E). Consistent with the fact that RIPK1 kinase activity is required for necroptosis induction (*36*), the selective RIPK1 inhibitor GSK’963 suppressed the TE- and TBE-mediated killing (Fig. 2B, 2C, 2D and 2E). In addition, D377A mutant MDFs were also significantly more sensitive than WT MDFs to TNF treatment following expression of CrmA (*37*), a Cowpox virus-encoded Caspase-8 inhibitor, via a doxycycline induced construct (Fig. 2F and S3B). Necroptosis can also be triggered by IFNg-induced ZBP1 upregulation, in combination with Caspase-8 inhibition (*25*). Interestingly, cFLIP mutant cells were also more sensitive than WT cells to IFNg and emricasan (IE)-induced necroptosis, in a TNF-independent manner (Fig. 2G and S3C). cFLIP mutant cells were also more sensitive to TNF and birinapant (TB)- and TNF and cycloheximide (TC)-induced apoptosis (Fig. 2H and 2I, 2J and S3D). TB-induced cell death was also RIPK1 kinase-dependent, since the GSK’963 inhibitor could abolish cell death both in WT and mutant cells (Fig. 2H, 2I and 2J). Finally, the genotoxic drugs Gemcitabine, Paclitaxel and Doxorubicin, known to activate intrinsic apoptosis, killed WT and D377A mutant cells to the same extent (Fig. S3E, S3F, S3G and S3H), indicating the sensitization effect imparted by the cFLIP mutation is specific for the TNF signalling pathway. Altogether, these data indicate that cFLIP cleavage specifically limits TNF-induced apoptosis and necroptosis in multiple cell types, and that it plays no role in cell death processes induced by genotoxins.

**Fig. 2.**
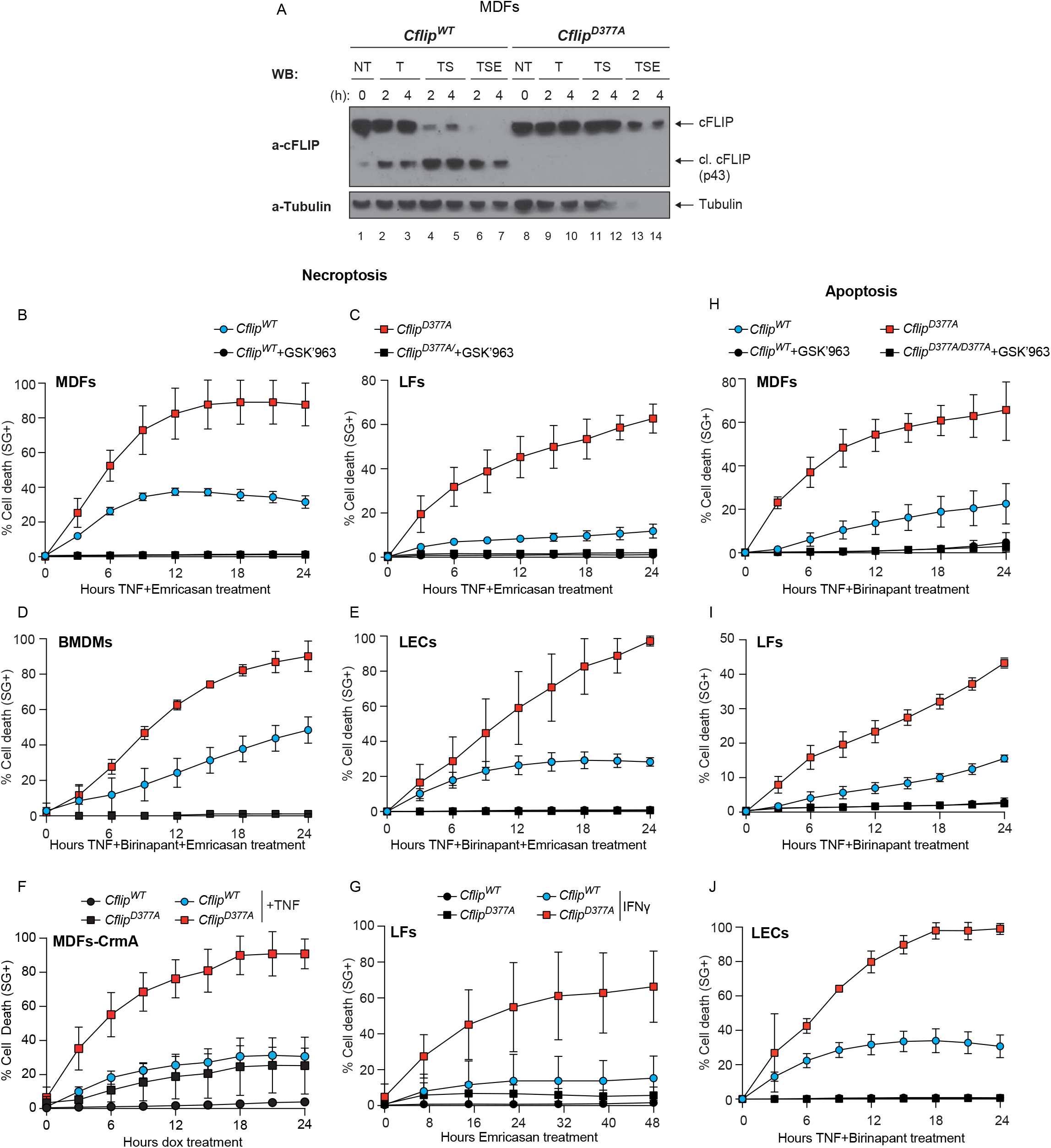
Cleavage resistant cFLIP mutant sensitizes to TNF-induced necroptosis and apoptosis. (**A**) *Cflip^WT^* and *Cflip*^*D377A*^ MDFs were treated as indicated (TNF 100 ng/ml, emricasan 1 μg/ml and birinapant 250 nM) and cell lysates were analyzed by immunoblotting using the indicated specific antibodies (n=2). (**B**) WT and D377A mutant MDFs, (**C**) lung fibroblasts (LFs), (**D**) bone marrow derived macrophages (BMDMs) and (**E**) lung endothelial cells (LECs) were treated with TNF and emricasan or TNF, the Smac mimetic birinapant and emricasan, in the presence or not of the RIPK1 specific inhibitor GSK’963 (100 nM) and cell death was measured over time by calculating the percentage of Sytox Green positive cells. For MDFs 1 ng/ml TNF and 1 μM emricasan (n=6), for LFs 10 ng/ml TNF and 1 μM emricasan (n=4), for BMDMS 10 ng/ml TNF, 250 nM birinapant and 1 μM emricasan (n=5) and for LECs 10 ng/ml TNF, 250 nM birinapant and 1 μM emricasan (n=3). (**F**) MDFs cFLIP WT and D377A stably expressing a doxycycline-inducible HA-tagged CrmA construct were treated for 48 hours with doxycycline (1μg/ml) and then treated or not with TNF (100 ng/ml). Cell death was measured as in B-E (n=3). (**G**) WT and D377A mutant LFs were pretreated with IFNγ for 18 hours and then subjected to emricasan (1μm) treatment. Cell death was measured as in B-E (n=3). (**H**) WT and D377A mutant MDFs, (**I**) LFs and (J) LECs were treated with TNF and birinapant in the presence or not of the RIPK1 specific inhibitor GSK’963 (100 nM) and cell death was measured over time by calculating the percentage of Sytox Green positive cells. For MDFs, LFs and LECs 10 ng/ml TNF and 250nM birinapant, n=4.

### The D377A mutation enhances complex-II formation, independently of NF-kB

TNF-induced necroptosis is driven by the formation of the RIPK1-kinase activity-dependent complex-II and the consequent phosphorylation of RIPK3 and MLKL (*23, 36*). To ascertain how the abrogation of cFLIP cleavage sensitizes cells to necroptosis, we treated MDFs and BMDMs with TE or TBE, respectively, and immunoprecipitated complex-II via FADD. The subsequent immunoblotting analysis revealed increased association of RIPK1, RIPK3, Caspase-8 and cFLIP with FADD in the *Cflip*^*D377A*^ cells (Fig. 3A and 3B). In addition, we observed significantly higher levels of phosphorylated RIPK1, RIPK3 and MLKL in cFLIP mutant cells treated with TE or TSE, IE or expressing CrmA (Fig. 3C, 3D and S4A and 4B). In addition to phosphorylation, RIPK1 and MLKL also undergo ubiquitin modification that promotes their killing activity during necroptosis. Consistent with *Cflip*^*D377A*^ cells being more sensitive to necroptosis, we observed dramatically higher ubiquitination of RIPK1, MLKL and cFLIP in the mutant cells compared to WT cells, upon TE treatment (Fig. 3E and S4C). TNF-induced apoptosis is mediated by Caspase-8 activation, by means of auto-proteolytic cleavage, that in turn activates the executioner caspases, Caspase-3 and 7(*38*). Consistent with the observation that *Cflip*^*D377A*^ cells are more sensitive to apoptosis, we observed increased cleaved Caspase-8 and cleaved Caspase-3 in cFLIP mutant cells treated with TB or TC (Fig. 3F and 3G). Finally, the enhanced sensitivity to cell death imparted by the cFLIP mutation is not due to defects in activation of NF-kB or MAPKs, since these signalling pathways were not affected by the D377A mutation (Fig. S4D and S4E). Collectively, these data demonstrate that the D377A mutation promotes complex-II formation and its ability to induce MLKL-dependent necroptosis and Caspase-3-dependent apoptosis.

**Fig. 3.**
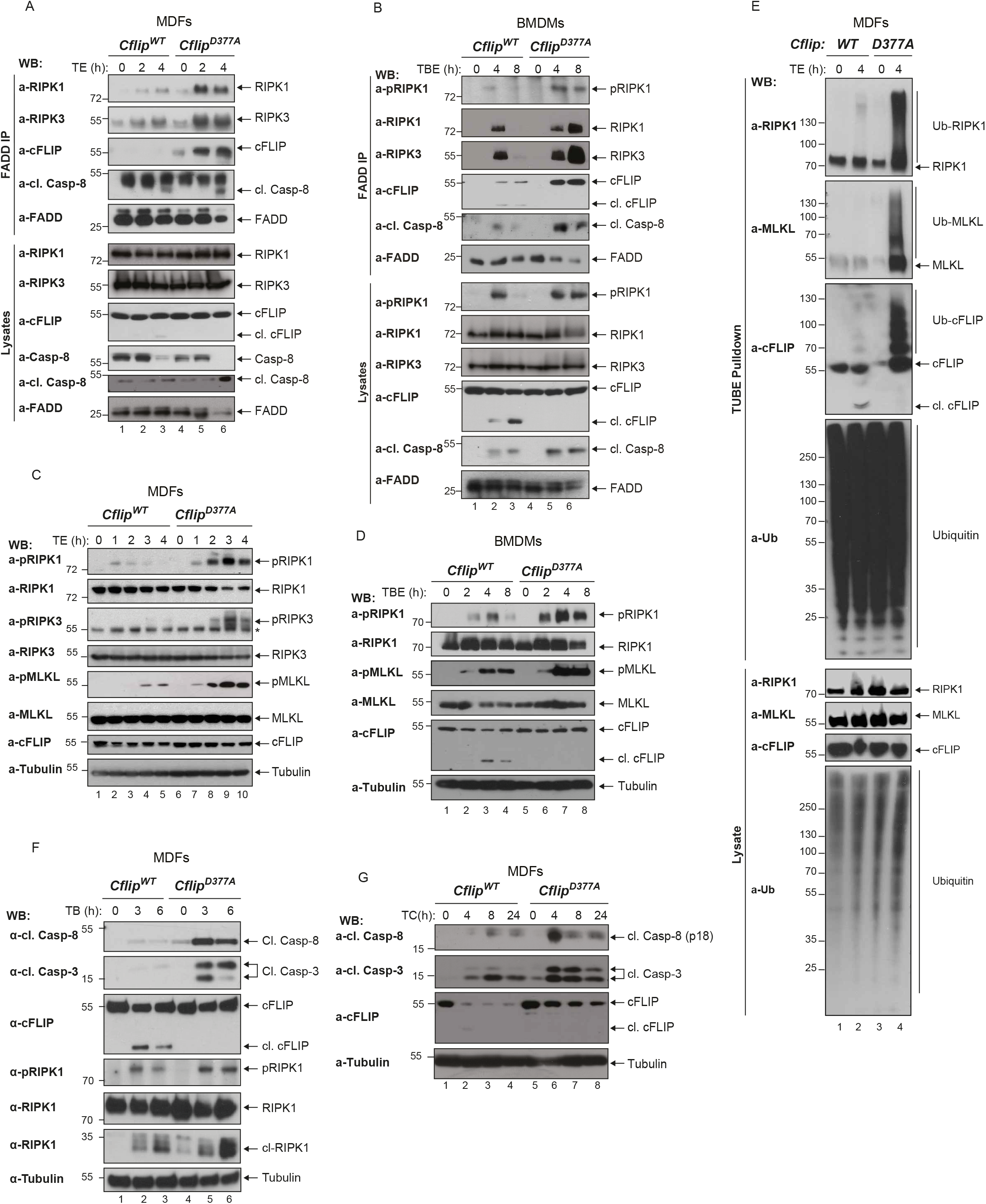
The D377A mutation enhances complex-II formation, independently of NF-kB. (**A**) WT and D377A-mutant MDFs and (**B**) BMDMs were treated with TNF (1 ng/ml) and emricasan (1 μg/ml) and TNF (1 ng/ml), birinapant (250 nM) and emricasan (1 μg/ml), respectively, for the indicated time points. Cell lysates were subjected to immunoprecipitation using a FADD specific antibody. Immunocomplexes and cellular lysates were then analyzed by immunoblotting using the indicated specific antibodies (n=3). (**C**) MDFs and (**D**) BMDMs were treated as in A and B for the indicated time points. Cell lysates were analyzed by immunoblotting using the indicated specific antibodies (n=2). (**E**) MDFs were treated with TNF (100 ng/ml) and emricasan (1 μg/ml) and cell lysates were subjected to TUBE pull-down, followed by immunoblotting analysis with the indicated specific antibodies (n=3). (**F**) WT and D377A-mutant MDFs were treated with TNF (10 ng/ml) and birinapant (250 nM) or (**G**) TNF (10 ng/ml) and cycloheximide (1 μg/ml) for the indicated time points and cell lysates were immunoblotted using the indicated specific antibodies (n=2).

### cFLIP cleavage arrests complex-II formation

To mechanistically understand how cFLIP cleavage limits TNF-induced complex-II formation and cell death, we performed size-exclusion chromatography. As expected, in WT cells treated with TBE, a small portion of RIPK1, RIPK3, cFLIP and cleaved Caspase-8 co-eluted with FADD in fractions 17-21, that contain high molecular weight complexes of apparent molecular mass of ~2 MDa. Immunoprecipitation of FADD from pooled column fractions (17-21) further confirmed that these proteins are together in the same complex (Fig. S5A). When gel filtration profiles of lysates from *Cflip^WT^* and *Cflip*^*D377A*^ MDFs treated with TBE were compared, we observed increased abundance of RIPK1, RIPK3, cleaved Caspase-8 and cFLIP in fractions 17-21, corresponding to the high molecular weight complex (Fig. 4A). However, the D377A mutation did not further increase the mass of the high molecular weight complex, since in both *Cflip^WT^* and *Cflip*^*D377A*^ MDFs complex-II components co-eluted in the same fractions (17-21). This corroborated the results obtained with FADD immunoprecipitation (Fig. 3A and 3B) and indicated that cleavage of cFLIP at position D377 limits the extent of complex-II formation. We next reasoned that the p12 domain of cFLIP, which is C-terminal to the D377A, might be important for the assembly and stability of complex-II. This fragment contains a residue, glutamine (Q) 469, that was reported to be important for cFLIP heterodimerization with Caspase-8(*31*) (Fig. 4B). Thus, cFLIP cleavage could regulate formation of the cFLIP/Caspase-8 heterodimers and, as a consequence, the assembly, stabilization and killing potential of complex-II. To interrogate this possibility, we reconstituted *Cflip^-/-^* MDFs with different cFLIP mutant constructs (Fig. 4B). If our hypothesis were correct, the Q469D mutation would abrogate the pro-cell death effects of the D377A mutation. Very interestingly, while D377A-reconstituted cells were significantly more sensitive to TE-induced cell death than WT-reconstituted cells, the Q469D mutation reverted the sensitizing effect of the D377A mutation (Fig. 4C). Indeed, the levels of TE-induced cell death in D377A/Q469D-reconstituted cells were comparable to those observed in WT-reconstituted cells. Consistent with this, phosphorylated MLKL levels in WT- and D377A/Q469D-reconstituted-cells were reduced when compared to D377A-reconstituted cells (Fig. 4D). Altogether, these findings indicate that the cell death sensitizing ability of the D377A depends on the Q469. This supports a model whereby cFLIP cleavage represents a molecular mechanism that controls the availability of a cFLIP/Caspase-8 interaction surface, the Q469, to limit the extent to which complex-II can form and mitigate cell death responses.

**Fig. 4.**
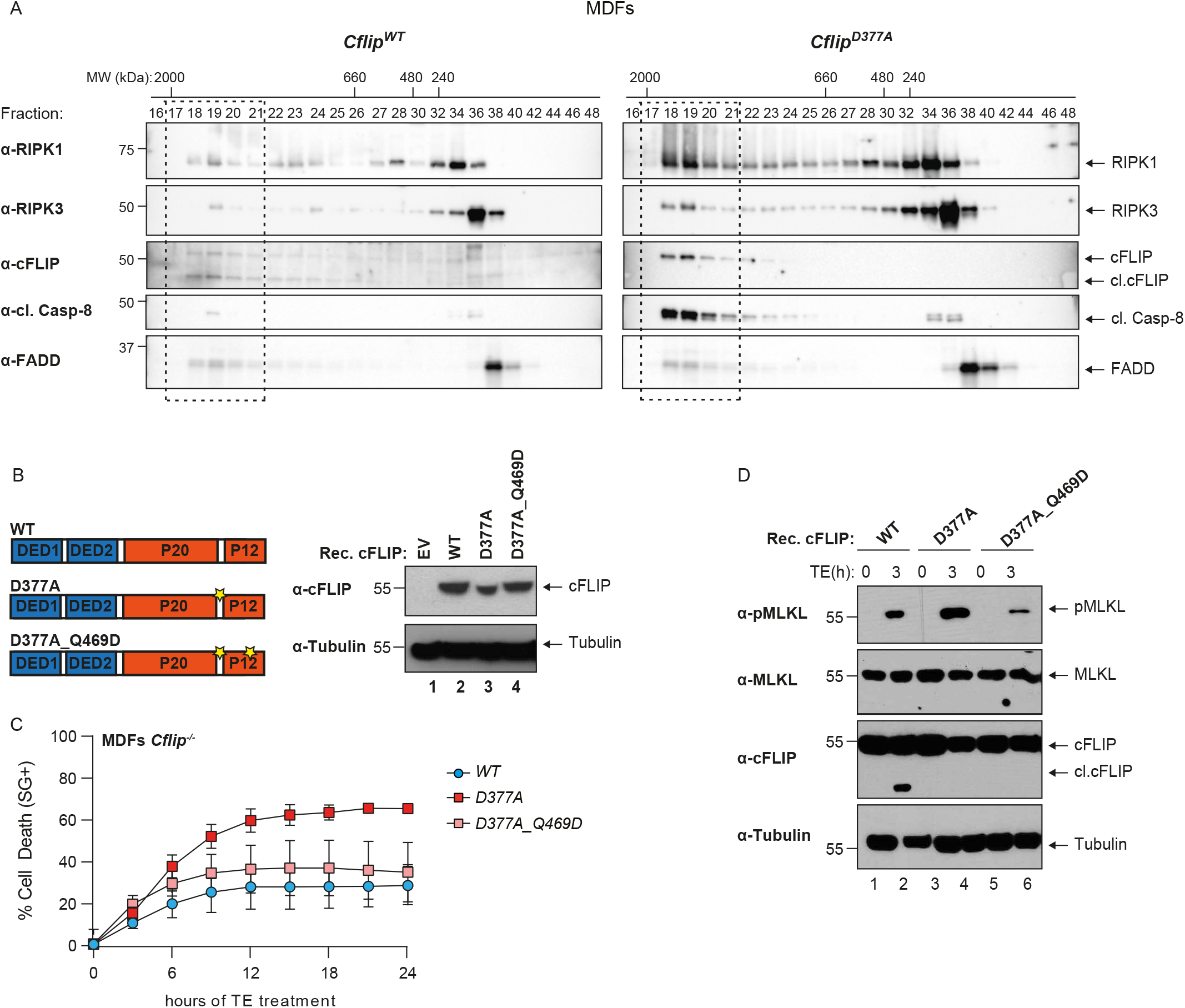
cFLIP cleavage counteracts complex-II formation. (**A**) *Cflip^WT^* and *Cflip*^*D377A*^ MDFs were treated with TNF (10 ng/ml), Smac mimetic (250 nM) and emricasan (1 μg/ml) for 4 hours and lysates were separated on a Superose 6 size-exclusion column. Aliquots from each fraction were retained and analyzed by immunoblotting with the indicated specific antibodies. (**B**) Cartoon depicting cFLIP domain composition and position of the D377 and Q469 residues (left part). Immunoblotting analysis of *Cflip^-/-^* MDFs reconstituted with the indicated cFLIP constructs via lentiviral infection, or with an empty lentivirus (EV). cFLIP- and Tubulin-specific antibodies were used (right panel). (**C**) *Cflip^-/-^* MDFs reconstituted as in B were treated with TNF (1 ng/ml) emricasan (1 μg/ml) and cell death was measured over time by calculating the percentage of Sytox Green positive cells (n=3). (**D**) MDFs *Cflip^-/-^* MDFs reconstituted as in C and treated as in D for 4 hours. Cell lysates were analyzed by immunoblotting with the indicated specific antibodies.

### Abrogation of cFLIP cleavage exacerbates the phenotype of *Sharpin* mutant mice

Next, we sought to investigate in a genetic model of TNF-induced cell death-mediated tissue damage whether the cFLIP cleavage had cell death-limiting functions. Sharpin deficiency in mice causes a TNF-mediated, cell death-dependent multiorgan inflammation, referred to as chronic proliferative dermatitis mice (cpdm), characterized by skin dermatitis, lung and liver inflammation, loss of marginal zones in the spleen and loss of Peyer’s patches (*39–41*). Since cleavage at D377 limits TNF-induced complex-II-mediated cell death, we set up to investigate whether the loss of cFLIP cleavage would exacerbate the effects of Sharpin deletion *in vivo*. For this purpose, we generated *Cflip^D377A/D377A^Sharpin^cpdm/cpdm^* double mutant mice (*Cflip^D377A^Sharpin^cpdm^*). The *Cflip*^*D377A*^*Sharpin*^*cpdm*^ were born at the expected mendelian ratio, yet they were runted and displayed significantly lower body weight than *Sharpin^cpdm/cpdm^* mice (*Sharpin^cpdm^*) (Fig. 5A). Although *Sharpin*^*cpdm*^ mice exhibited splenomegaly, the D377A mutation did not cause further spleen enlargement (Fig. S6A). However, we observed a complete loss of marginal zones in *Cflip^D377A^Sharpin^cpdm^* mice, while *Sharpin*^*cpdm*^ exhibited only minor spleen architecture alterations at 3 and 7 weeks of age (Fig. 5B). Consistently with this finding, we detected a significantly higher number of TUNEL-positive cells in the spleen of *Cflip^D377A^Sharpin^cpdm^* mice compared to *Sharpin*^*cpdm*^ and control mice (Fig. 5C). *Sharpin*^*cpdm*^ developed skin lesion between 15 and 20 weeks after birth (Fig. S6B). When *Cflip^D377A^Sharpin^cpdm^* reached the termination criteria, around week 10 after birth, no sign of dermatitis was detected. However, skin samples of 3-weeks-old *Cflip^D377A^Sharpin^cpdm^* mice revealed a significantly increased amount of TUNEL-positive cells in the skin compared to *Sharpin*^*cpdm*^ and control mice (Fig. 5D). *Sharpin*^*cpdm*^ mice are also characterized by the loss of Peyer’s patches (PP) (Fig. S6C). Despite the loss of PPs, histological analysis revealed that intestines of 7-weeks-old *Sharpin*^*cpdm*^ mice were normal and largely comparable to those of control mice (Fig. 5E). On the contrary, intestines of *Cflip^D377A^Sharpin^cpdm^* mice exhibited signs of damage, inflammation and cell death, both in the small and large intestine, as shown by the histology score (Fig. 5E), that correlates with a significant increase in the amount of TUNEL-positive cells (Fig. 5F). Altogether, these findings indicate that the D377A mutation accelerates cell death induced by Sharpin deletion in the spleen, skin and intestine. Therefore, we conclude that the abrogation of cFLIP cleavage enhances the cell death-inducing potential of Sharpin deletion *in vivo*.

**Fig. 5.**
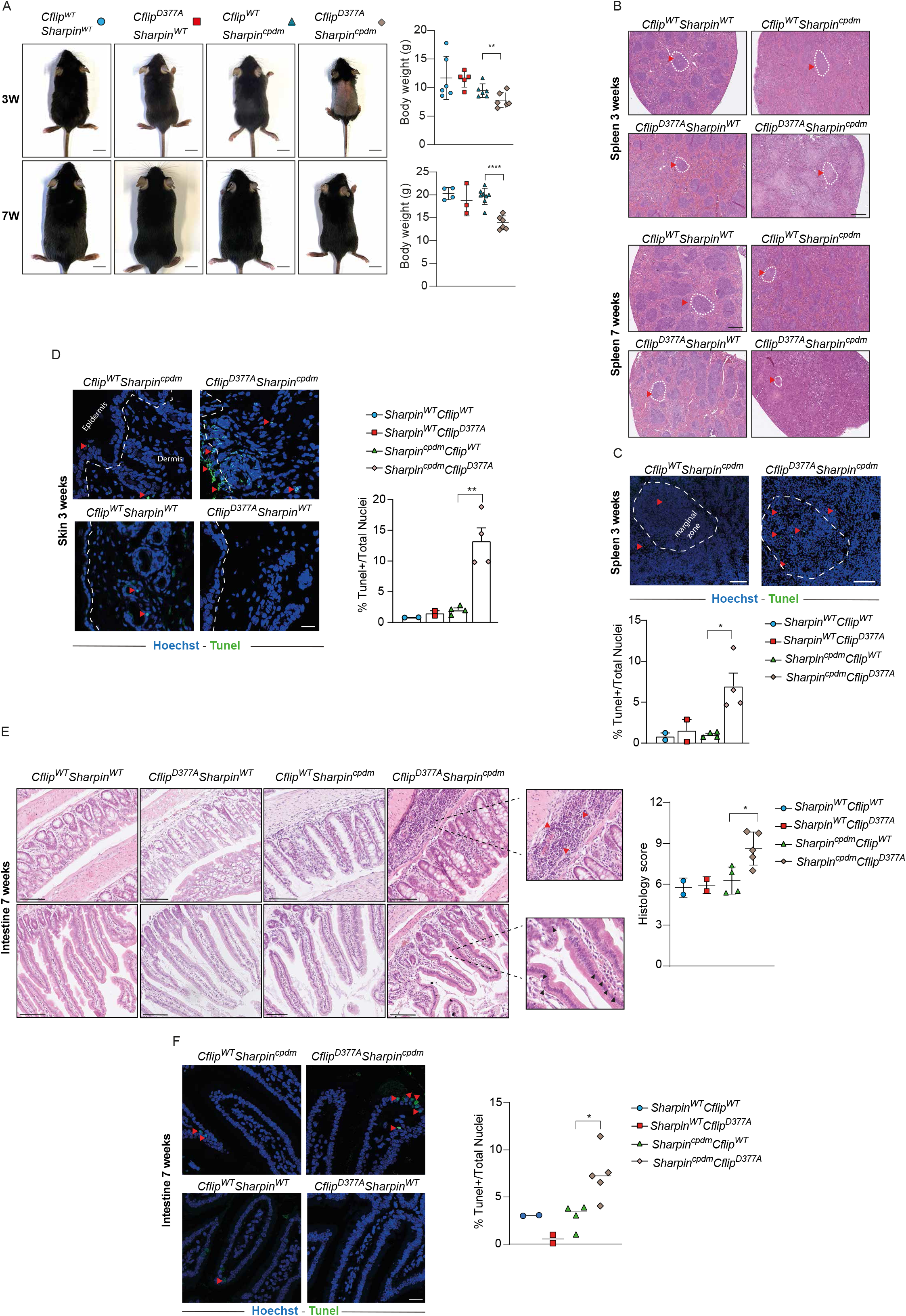
Abrogation of cFLIP cleavage exacerbates the phenotype of *Sharpin* mutant mice. (**A**) Pictures (left) and body weight (right) of 3 and 7 weeks old *Cflip^WT^Sharpin^WT^, *Cflip*^*D377A*^*Sharpin^WT^*, Cflip^WT^Sharpin^cpdm^* and *Cflip*^*D377A*^*Sharpin^cpdm^* mice. Each symbol corresponds to one mouse. Scale bar 1 cm. Data are presented as mean ± SD, **p<0,01 (**B**) Spleen sections of 3- and 7-weeks-old mice of the indicated genotypes (as in A). Scale bar 500 μm. Dotted with circles and red arrowheads indicate marginal zones. (**C**) TUNEL staining of spleen sections of 3 weeks old mice of the indicted genotypes (upper part) and the relative quantification (lower part) expressed as percentage of TUNEL positive cells over the total number of cells. Hoechst stains nuclei. Red arrowheads indicate TUNEL positive cells. Data are presented as mean ± SD, *p<0,05. Scale bar 100 μm. (**D**) TUNEL staining of skin sections of 3 weeks old mice of the indicated genotypes (left) with the relative quantification (right) expressed as percentage of TUNEL positive cells over the total number of cells. Scale bar 20 μm. Data are presented as mean ± SD, **p<0,01 (**E**) Large (top row) and small (lower row) intestine sections of 7 weeks old mice of the indicated genotypes with the histology score relative to the whole intestine. Each symbol represents one mouse. Data are presented as mean ± SD, *p<0,05. Scale bar 100μm. Red and dark arrowheads indicate areas of inflammation and dead cells, respectively. (**F**) TUNEL staining of small intestine sections of 7 weeks old mice of the indicated genotypes (left) with the relative quantification (right) expressed as percentage of TUNEL positive cells over the total number of cells. Scale bar 20 μm. Red arrowheads indicate TUNEL positive cells. Data are presented as mean ± SD, *p<0,05.

### cFLIP^D377A^ mice exhibit impaired skin wound healing

To further validate the role of cFLIP cleavage in protecting tissues from cell death-induced damage, we used a model of mouse full-thickness excisional skin injury. Since Caspase-8 was reported to play a crucial role in the wound healing response (*42*), we wondered whether Caspase-8-mediated cFLIP cleavage could contribute to wound closure. The wound closure is characterized by the formation of the granulation tissue in the damaged area that is composed of different cell types whose concerted action promotes wound closure (*43, 44*) (Fig. 6A). Endothelial cells, myofibroblasts and macrophages are the main components of the granulation tissue (*44*). Interestingly, histomorphological analysis indicated a significant delay in the dynamics of wound closure in *Cflip*^*D377A*^ mice as compared to WT mice, as shown by reduced scar formation (Fig. 6B) and reduced granulation tissue area at days 4, 7 and 14 after injury (Fig. 6C and 6D). Consistent with a reduction in the granulation tissue, *Cflip*^*D377A*^ mice exhibited significantly higher levels of TUNEL-positive cells (Fig. 6E and 6F), reduced wound vascularization (CD31 staining) and less myofibroblasts (α-SMA staining) at day 7 after injury (Fig. 6G and 6H). Thus, impairing cFLIP cleavage causes an increased cell death response that interferes with the wound healing response. We therefore conclude that cFLIP cleavage controls the extent of cell death occurring after wound injury to favor an optimal healing process.

**Fig. 6.**
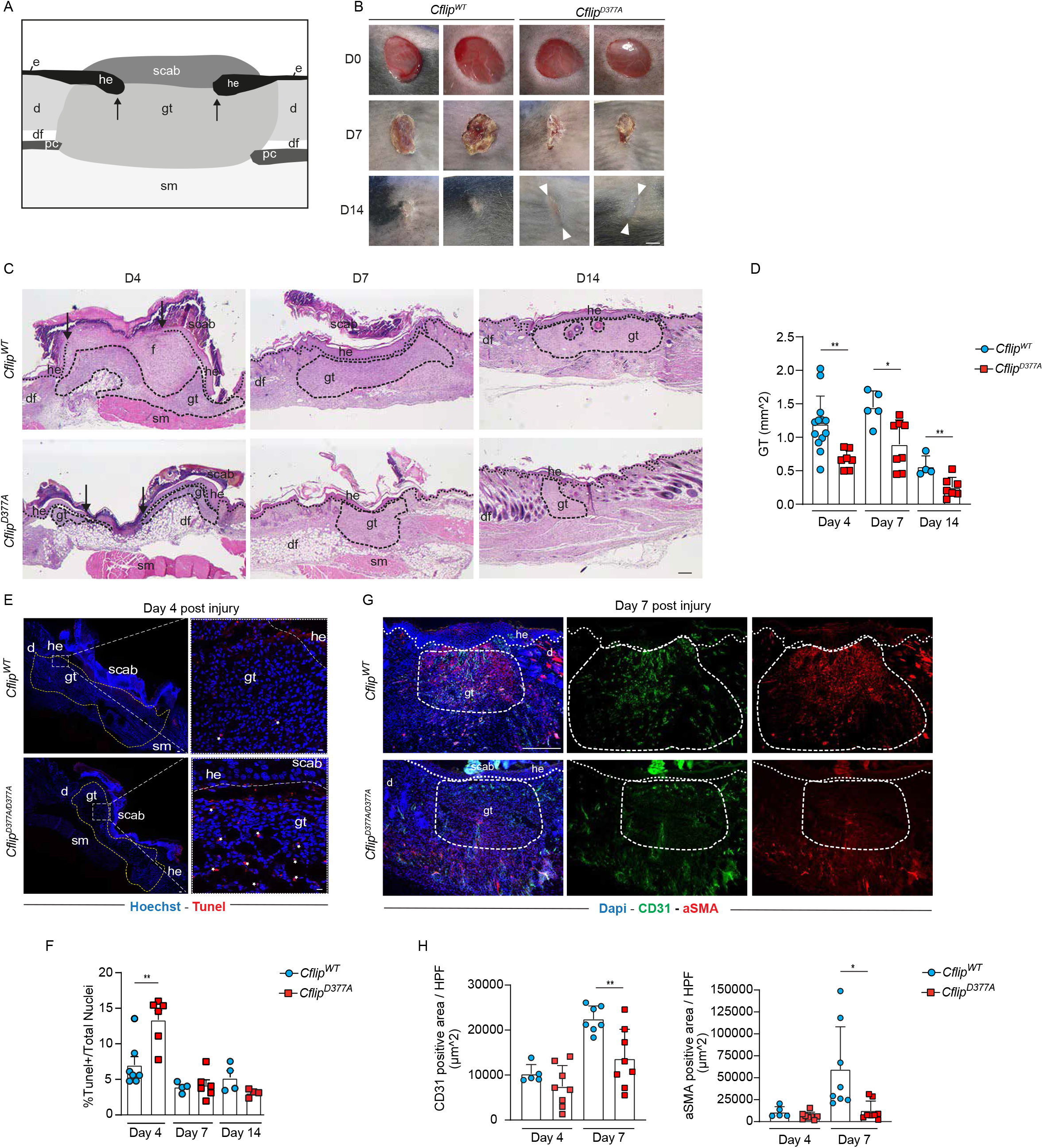
*cFLIPD^377A^* mice exhibit impaired skin wound healing. (**A**) Cartoon representing the histology of a skin wound at around 4 dpi, where the following skin areas are marked: d, dermis; df, dermal fat tissue; e, epidermis; gt, granulation tissue; he, hyperproliferative epithelium; pc, panniculus carnosus; sm, skeletal muscle. (**B**) Macroscopic pictures of wounds at 0, 7 and 14 days post injury (dpi) in *Cflip^WT^* and *Cflip^D377A^* mice. White arrows indicate scar tissue. Arrows indicate the tips of the epithelial tongues. Scale bar 2 mm. (**C**) Representative H&E-stained wound sections of *Cflip^WT^* (upper panels) and *Cflip*^*D377A*^ mice (lower panel) at 4, 7 and 14 dpi. Scale bar 200 μm. (**D**) Quantitative analysis of granulation tissue (gt) area of sections in C at 4, 7, and 14 dpi (n = 4-12 total wounds per genotype per dpi). Scale bar 200 μm. (**E**) Representative TUNEL-stained wound sections at 4 dpi of *Cflip^WT^* mice and *Cflip^D377A^* mice (white stars indicate TUNEL positive cells). Nuclei were stained with Hoechst (blue). Scale bars 50μm in left images and 20μm in right images. (**F**) Percentage of total TUNEL positive cells as represented in E at 4, 7, and 14 dpi (n= 4-7 total wounds per genotype per dpi). (**G**) Representative CD31 (green) and αSMA (red) immunostainings at day 7 dpi of wound sections of *Cflip^WT^* and *Cflip^D377A^* mice. Scale bar 200μm. (**H**) Percentage of total CD31 and αSMA positive area per hpf (high power field) in wound sections as represented in G, at 4, and 7dpi (n = 5-8 total wounds per genotype per dpi).

### cFLIP^D377A^ mice are more sensitive to SARS-CoV-induced lethality

Different types of viruses, including coronaviruses, can induce cell death by triggering Caspase-8-dependent apoptosis or ZBP1/RIPK3 dependent necroptosis, directly in the infected cells or as a consequence of the hyperinflammatory state created by the immune system (*45–47*). Our observation that the D377A mutation sensitizes cells to both apoptosis and necroptosis, prompted us to investigate the biological role of cFLIP cleavage in a model of SARS-coronavirus (SARS-CoV) infection. We therefore infected WT and *Cflip*^*D377A*^ mice with a sub-lethal dose of a mouse-adapted strain of SARS-CoV (*48*) (MA15), known to induce the production of TNF and other cytokines in the lung of infected mice (*49, 50*). Strikingly, *Cflip*^*D377A*^ mice were dramatically more sensitive than WT mice to the lethal effect of the virus (Fig. 7A). However, we could not detect any difference in terms of viral titer between the two genotypes (Fig. 7B). We therefore reasoned that cFLIP cleavage is not involved in controlling viral replication. In addition, the levels of pro-inflammatory cytokines and chemokines between WT and mutant mice were largely comparable (Fig. 7C). When examining lung sections of infected mice, we detected a significantly higher number of TUNEL-positive cells in the lungs of infected *Cflip*^*D377A*^ mice compared to infected WT infected mice, both in the bronchioles and alveoli (Fig. 7D). In addition, co-staining with TUNEL and CC10, a marker of club cells which are present in the respiratory bronchioles, revealed a higher number of TUNEL-positive club cells in *Cflip*^*D377A*^ mice, indicating increased lung tissue damage (Fig. 7E). To explain the augmented cell death levels observed in the lungs of *Cflip*^*D377A*^ mice we considered the possibility that the cFLIP-mutant lung cells exhibited higher sensitivity to the cytokines produced during the viral response. TNF and INFγ are among the main produced cytokines during SARS-CoV infection. Therefore, we pretreated WT and D377A mutant lung fibroblasts (LFs) with IFNγ prior to TNF treatment and found that cFLIP mutant LFs are significantly more sensitive than their WT counterparts to IFNγ/TNF-induced cell death (Fig. 7F). Immunoblotting analysis revealed that mutant LFs underwent necroptosis, as shown by the higher levels of phosphorylated MLKL (Fig. 7G). This suggested that cFLIP cleavage can control the killing activity of a cytotoxic complex induced by TNF and IFNγ.

**Fig. 7.**
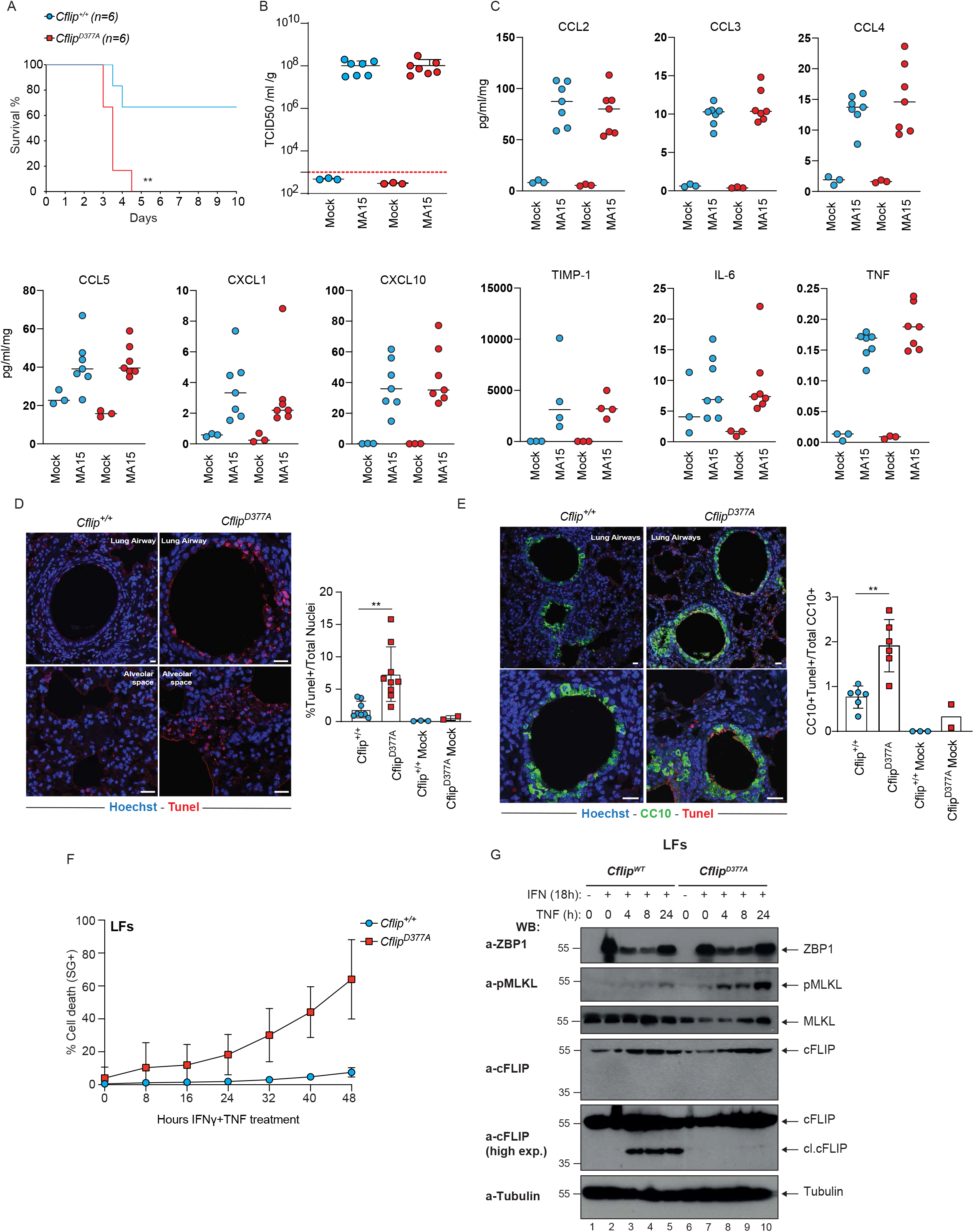
cFLIP^D377A^ mice are more sensitive to SARS-CoV-induced lethality. (**A**) Survival curve of *Cflip^WT^* and *Cflip^D377A^* mice infected with SARS-Cov MA15 virus (n=6). Survival curves were compared using log-rank Mantel–Cox test, **p<0,01. Viral titer (**B**) and cytokine levels (**C)** from lungs of *Cflip^WT^* and *Cflip*^*D377A*^ mice 3 dpi (n=3 for mock infection and n=7 for MA15 infection). (**D**) Representative TUNEL-stained lung sections of *Cflip^WT^* mice and *Cflip*^*D377A*^ mice 3 dpi (left). Nuclei were stained with Hoechst (blue). Scale bars 10μm (upper left image), and 20μm (upper right and bottom images). Percentage of total TUNEL positive cells (right) (n=8 for *Cflip^WT^* infected lungs, n=10 *Cflip^D377A^* infected lungs and n=2-3 for mock infection). Data are presented as mean ± SD, **p<0,01. (**E**) Representative CC10 (green)/TUNEL (red) double stained lung sections of *Cflip^WT^* mice and *Cflip^D377A^* mice 3 dpi (left). Nuclei were stained with Hoechst (blue). Scale bars 10μm (upper images) and 20μm (bottom images). Percentage of total CC10/TUNEL double positive cells (right). (n=6 for *Cflip^WT^* infected lungs, n=6 *Cflip*^*D377A*^ infected lungs and n=2-3 for mock infection). Data are presented as mean ± SD, **p<0,01. (**F**) *Cflip^WT^* and *Cflip*^*D377A*^ lung fibroblasts (LFs) where pre-treated with IFNγ (100nM) for 18 hours and then treated with TNF (100 ng/mL) for additional 24 hours. Cell death was measured over time by calculating the percentage of Sytox Green positive cells (n=3). (**G**) Total lysates of LFs treated like in F for the indicated times points were immunoblotted with the indicated specific antibodies (n=2).

Altogether, these findings show that cFLIP cleavage represents a mechanism that protects mice from SARS-CoV-induced lethality by limiting the extent of cytokine-induced cell death occurring in the lungs.

## Discussion

cFLIP is an essential, non-redundant cell death suppressor required to maintain tissue integrity(*51–56*). Its cell death inhibitory functions are due to the fact that it heterodimerizes with Caspase-8 and modulates its activity. Apart from being a modulator of Caspase-8 activity, cFLIP is also a Caspase-8 substrate, cleaved at position D377(*32*). However, despite earlier studies on cFLIP proteolysis at D377(*31, 33, 34, 57*), the biological relevance of this cleavage events remained to be elucidated. Here we generated mutant mice carrying a non-cleavable cFLIP version, the *Cflip*^*D377A*^, which were born at the expected mendelian ratio. It was previously shown that cFLIP is required for Caspase-8-mediated suppression of necroptosis. Our results indicate that the cleavage of cFLIP is dispensable for the ability of Caspase-8 to suppress necroptosis. When we analyzed different D377A mutant mouse cells, including bone marrow derived macrophages and lung endothelial cells, we found that the abrogation of cFLIP cleavage sensitizes them to TNF-mediated apoptosis and necroptosis (induced by TNF and Smac mimetic and TNF and emricasan, respectively), by enhancing complex-II formation. Interestingly, the D377A sensitizes cells not only to RIPK1 dependent cells death (apoptosis and necroptosis) but also to RIPK1-independent apoptosis (TNF and cycloheximide) and ZBP1-dependent necroptosis (IFNγ and Emricasan). Of note, in a previous study, a mutant mouse expressing non-cleavable cFLIP was generated, where both D377 and the nearby D371 were mutated (*35*). Here, we show that the D377A mutation is sufficient to render cFLIP cleavage resistant.

*In vivo* the *Cflip*^*D377A*^ mice were more susceptible to SARS-CoV-mediated lethality than WT mice, and this correlated with significantly higher levels of lung epithelial cell death, while no difference in viral titer or cytokine production was observed in lung extracts. Therefore, we envisage a scenario whereby the D377A mutation sensitizes lung cells to cell death induced by the overproduction of cytokines following lung infection, referred to as cytokine storm, that include TNF and IFNγ. Therefore, the increased cell death observed in the lungs of mutant mice would not be the trigger but the consequence of the cytokine storm and the link between this last and the more severe lung pathology. Consistent with this scenario, we found that lung fibroblasts from the *Cflip*^*D377A*^ mouse are significantly more sensitive than WT cells to TNF/IFNγ-induced cell death. Since IFNγ induces the upregulation of ZBP1 that can in turn trigger the assembly of RIPK1/RIPK3/Caspase-8/cFLIP complex (*58*), we speculate that cFLIP cleavage can regulate the activity of this ZBP1-mediated cytotoxic complex. In addition, *Cflip*^*D377A*^ mice exhibited impaired wound healing following skin excision, as a consequence of increased cell death observed in the granulation tissue. Again, this would be consistent with a model whereby the D377A mutation renders granulation tissue cells more sensitive to death induced by the cytokines, including TNF, produced during the wound healing processes, likely by macrophages. To further provide understanding of the biological role of D377A mutation in vivo, we crossed the *Sharpin*^*cpdm*^ mice, that carry an inactivating mutation on *Sharpin* that causes TNF-dependent cell death-mediated systemic inflammatory syndrome (*41*), with the *Cflip*^*D377A*^ mouse. The resulting *Sharpin^cpdm^Cflip^D377A^* mice were runted and showed increased cell death levels in intestine and skin in the first weeks after birth compared to the *Sharpin*^*cpdm*^ mice. However, we could not determine whether the *Sharpin^cpdm^Cflip^D377A^* mice would develop skin dermatitis earlier than the *Sharpin*^*cpdm*^ mice or lethal intestinal inflammation, since they had to be terminated for ethic reasons around 10 weeks of age. At the mechanistic level we were able to show that the cleavage of cFLIP controls the extent of complex-II formation by regulating cFLIP/Caspase-8 heterodimerization via Q469. Two pieces of evidence support this model: i) in the D377A mutant cells stimulated with TNF, birinapant and emricasan, there is a higher abundance of complex-II components co-eluting in the 2 ~MDa gel filtration fractions. This suggests that the D377A mutation favors the assembly of complex-II. ii) the Q469D mutation abrogates the cell death sensitizing effects of the D377A mutation. This indicates that in the absence of cFLIP cleavage, Q469 favors complex-II assembly and stability most probably by promoting cFLIP/Caspase-8 heterodimerization. Caspase-8 can in turn recruit more FADD molecules, via DED-mediated interaction, that in turn can recruit more RIPK1 molecules, via DD-mediated interaction. Therefore, the Caspase-8-mediated cleavage of cFLIP at position D377 represents a mechanism that counterbalances complex-II formation to keep its killing activity in check. Previous studies have reported conflicting results regarding the role of cFLIP cleavage on Caspase-8 activity, mainly using cell-free system assays containing recombinant or hybrid cFLIP and Caspase-8 (*31, 33, 34*). Such systems do not recapitulate the complexity of cell death-inducing platforms, such as TNF-induced complex-II, that need to be assembled in order to activate Caspase-8. Therefore, in the light of our findings, we believe cFLIP cleavage does not directly impact on Caspase-8 activity per se, but rather acts indirectly, by regulating the dynamics of a complex to which Caspase-8 is recruited and at which it is activated.

The ability of tissues to overcome stress-induced damage lies also in their capacity to tightly control the amplitude of cell death responses, by modulating the formation and activity of cell death-inducing complexes. Indeed, if in the absence of cell death tissue repair programs cannot be initiated, excessive cell death can lead to hyperinflammatory responses that are in turn detrimental for the tissue. Different types of insults, such as viral and bacterial infections or mechanical tissue damage, are known to activate a cFLIP/Caspase-8/RIPK1-containing cell death complex. The results presented here reveal that the cleavage of cFLIP is a required regulatory module in the intricated process that controls the activity of this cytotoxic complex, thereby ensuring optimal cell death responses and the consequent activation of tissue repair programs.

## Material and Methods

### Mice generation

The *Cflip*^*D377A*^ mutant mice were generated by the MAGEC laboratory (WEHI) on a C57BL/6J background. To generate these mice, 20 ng/μl of Cas9 mRNA, 10 ng/μl of sgRNA (TTGATGGCCCATCTACCTCC) and 40 ng/μl of oligo donor (GCCAAAGCTCTTTTTTATTCAGAACTATGAGTCGTTAGGTAGCCAGTTGGAAGATAGCAGTCTGGAGGTAGCTGGGCCATCAATAAAAAATGTGGACTCTAAGCCCCTGCAACCCAGACACTGCACAACTCA) were injected into the cytoplasm of fertilized one-cell stage embryos generated from wild type C57BL/6J breeders. Twenty-four hours later, two-cell stage embryos were transferred into the uteri of pseudo-pregnant female mice. Viable offspring were genotyped by next-generation sequencing. Targeted animals were backcrossed twice to wild type C57BL/6J to eliminate off-target mutations

### Cell lines

Immortalized mouse dermal fibroblasts (MDFs) and lung fibroblasts (LFs), Platinum-E and HEK293T cells were cultured in Dulbecco’s modified Eagle’s Medium (DMEM) supplemented with 10% Fetal Bovine Serum (FBS), penicillin and streptomycin under 10% CO2. Lung endothelial cells were seeded on 0.1% gelatine-coated wells and cultured in a 1:1 mixture of EGM2 (PromoCell) and fully supplemented Dulbecco’s modified Eagle medium (DMEM) (Merck) (containing 20% FCS, 4 gr/L glucose, 2 mM glutamine, 1% penicillin–streptomycin (100 U/ml penicillin, 100 μg/ml streptomycin), sodium pyruvate 1% (1 mM), HEPES (20 mM) and 1% non-essential amino acids). Vero E6 cells were kindly supplied by Júlia Vergara from the Centro de Investigación en Sanidad Animal IRTA-CReSA (Barcelona, Spain). Spain). Vero E6 cells were grown in DMEM (Sigma) supplemented with 10 % FBS (Sigma), 2 mM Glutamax (Gibco), 100 U/ml penicillin (Sigma), 100 μg/ml streptomycin (Sigma), 0.25 μg/ml amphotericin B (Sigma), 1 % non-essential amino acids (Gibco) and 25 mM HEPES (4-(2-hydroxyethyl)-1-piperanzineethanesulfonic acid) (Biowest).

### Isolation of primary cells and immortalization

Mouse dermal fibroblasts (MDFs) were generated from the tail skin of 8-10 weeks old mice. Once separated from the tail, the skin was minced with a scalpel and digested in 3 ml of trypsin (Sigma T4049) for 1 hour at 37°C. 7 ml of DMEM were then added, the tube was manually shaken several times and the digested tail was then filtered through a 100 μM cell strainer. Cells were washed once with PBS and seeded in a 6 cm plate. 2-3 days later, they were infected with a SV40 Large T antigen expressing retrovirus for immortalization. Mouse lung fibroblasts (LFs) were generated from whole lungs of 8-10 weeks old mice. Once separated from non-lung tissue, the lungs were cut in very small pieces and digested in 1.5 ml of Collagenase Type 2 (Worthington, 44N15307B) for one hour at 37°C and 500 rpm. One digested lung was seeded on two 15 cm dishes containing 20 ml of DMEM. After two days the medium was changed and cells were grown until confluence.

Lung endothelial cells were isolated from the lungs of 8-12 weeks old mice. Lungs were briefly washed in PBS, minced, and digested in 1 ml of 0.5% collagenase type II (Merck Millipore C2-22) for 45 min at 37°C. The cell solution was then filtered through a 70 μm cell strainer and separated with magnetic beads (mouse CD31, Miltenyi Biotec) according to the manufacturer’s protocol. CD31^+^ endothelial cells were seeded on gelatine-coated wells and cultured in the endothelial cell medium mentioned above. After the first passage, cells were resorted using the same magnetic beads. Isolated primary endothelial cells were analysed using the anti-CD31 antibody to distinguish endothelial cells from other cell types. To generate Bone Marrow Derived Macrophages (BMDMs), bone marrow cells from tibia and femur of 2-month-old mice were seeded in non-coated Petri dishes and cultured for 6 days in Dulbecco’s modified Eagle medium + 10% fetal bovine serum +20% (v/v) L929 mouse fibroblast conditioned medium. MDFs and LFs were immortalized using an SV40 Large T-Antigen-encoding retrovirus. Isolated cells were routinely tested negative for mycoplasma contamination by PCR.

### Reagents and antibodies

The following reagents were used in this study: Birinapant (MedChem Express, HY-16591), Cislplatin (X, X), Cyclohexemide (X, X), Doxorubicin (X, X), Doxycyclin (X, X), Emricasan (Absource Diagnostics, S7775-0005), Gemcitabine (X, X) GSK-963 (SeleckChem, S8642), IFNγ (Peprotech, 315-05-100), mouse TNFα (Enzo Life Sciences, ALX-522-009-C050), Paclitaxel (X, X), Sytox Green (Thermo Fischer Scientific, S7020). TRAIL and FAS-L were gifts from Prof. Dr. Henning Walczak. The following antibiotics were used: Blasticidine (Invivogen, ant-bl-1), Hygromycin-B (Invivogen, ant-hg-1/5), Neomycin (Invivogen, ant-gn-5), Puromycin (Invivogen, ant-pr-1). The following antibodies were used: Caspase-7 Rabbit Ab (Cell Signalling Technology, 12827), CC10 (Santa Cruz, sc365992), CD31 (BD Pharmingen, 550274), cFLIP Rabbit Ab (Cell Signalling Technology, 56343), Cleaved Caspase-3 Rabbit Ab (Cell Signalling Technology, 9661), Cleaved Caspase-8 Rabbit Ab (Cell Signalling Technology, 9429S), smooth muscle actin-Cy3 conjugated (Sigma-Aldrich, C6198), FADD Mouse Ab (Millipore, 05-486 / 1F7), FADD Rabbit Ab (abcam, ab124812), MLKL Rat Ab (EMD Millipore, MABC604), Phospho-MLKL Rabbit Ab (abcam, ab196436), Phospho-RIPK1 Rabbit Ab (Cell Signalling, 31122), Phospho-RIPK3 Rabbit Ab (Cell Signalling, 57220), RIPK1 Mouse Ab (BD Bioscience, 610459), RIPK1 Rabbit Ab (Cell Signalling, 3493), RIPK3 Rabbit Ab (ProSci Inc., 2283), Tubulin Mouse Ab (Thermo Fisher/Sigma, T9026 / 047M4789V), Ubiquitin Mouse Ab (Santa Cruz Biotechnology, SC-8017), ZBP1 Mouse Ab (Adipogene, AG-20B-0010-C100).

### SARS-CoV infection

SARS-CoV-1 MA15 was kindly provided by Sonia Zúñiga and Luis Enjuanes (CNB-CSIC, Madrid, Spain). For the virus titration in lungs, previously weighed portions were homogenised in 500 μl DMEM with a GentleMACS Dissociator (Miltenyi), centrifugation 1500 rpm x 5 min and the supernatant taken. Virus titration was determined by TCID50/ml assay performing serial dilutions and calculated using the Ramakrishan newly proposed method formula (*59*).

Mice were infected intranasally with 10^6^ TCID50/ml in a total volume of 40 μl PBS after isoflurane anesthesia. Mice were weighed daily and reached humanitarian endpoint with a 25 % weight loss. A clinical score was generated if needed, following different components: mouse appearance, level of consciousness, activity, response to stimuli, eye appearance, and frequency and quality of respiration; and mice reached humanitarian end point when the clinical score reached 21, the respiratory characteristics were higher than 3 or if the weight loss were greater than 25% of the initial weight. Animals were kept under standard conditions of temperature, humidity and light at the Research Center on Encephalopathies and Emerging Diseases of the University of Zaragoza. Animal experimentation was approved by the Animal Experimentation Ethics Committee of the University of Zaragoza (number: PI44/20).

### Excisional punch injury

Mice were anesthetized by intraperitoneal injection of 100 mg/kg body weight Ketavet (Pfizer) and 10 mg/kg bodyweight Rompun 2% (Bayer). The back skin was shaved using an electric shaver and disinfected with 70% ethanol. Full-thickness punch biopsies were created on the back using a standard biopsy puncher (Stiefel). For histological analysis, wounds were excised at different times after injury and processed following as described in Willenborg et al., 2012. The tissue was either fixed for 2 h in Roti Histofix or embedded in O.C.T. compound (Fisher Scientific) and stored at −80°C.

### Retroviral production and infection

Replication incompetent retroviral particles were generated in Platinum E cells. The cells were seeded in a 10-cm dish at 2.500.000 cells/dish in 10 ml culturing medium. The following day, 1 ml of Opti-MEM™ I Reduced Serum Medium containing 10 μg of plasmid of interest and 30 μl of 1 mg/ml polyethylenimine (PEI) were added to the medium and incubated overnight. The medium was changed after 18 h incubation. The supernatant containing retroviral particles was collected 2 days post-transfection. One day prior to the infection of target cells with retroviral particles, 80.000 cells/well were seed in 6-well plate. On the day of infection, 3 ml of supernatant-containing retroviral particles and 1 μg/ml polybrene were added to the respective wells of 6-well plate and kept for 72 h. To selectively expand the infected cells, 7-days selection was performed using the respective antibiotics depending on the resistance cassette contained in the retroviral plasmid, i.e Puromycin (4 μg/ml), Hygromycin (200 μg/ml), Blasticidine (6 μg/ml) and Neomycin (1 mg/ml).

### Cell culture, constructs and transfection

The SV40 Large T-Antigen and all the mouse cFLIP WT and mutant coding sequences (D371A, D377A, D371A/D377A, D377A/Q469D and □p12) were in the pBABE retroviral vectors, carrying either Puromycin or Hygromycin or Neomycin or Blasticidin resistance. *Cflip^-/-^* MDFs were generated from immortalized *Cflip^f/f^* MDFs infected with a Cre-expressing retrovirus, followed by antibiotic selection. All the reconstitutions of *Cflip^-/-^* MDFs were done by infection with retroviruses expressing the different cFLIP sequences, followed by antibiotic selection.

Mouse cFLIP WT was amplified from mouse cDNA and cloned into pBABE using BamHI and EcorRI restriction enzymes. All the cFLIP mutants were generated by site-directed mutagenesis and clones into pBABE using BamHI and EcorRI restriction enzymes.

### Complex-II purification

Cells were seeded in 10 cm dishes and treated as indicated using media containing mTNF (100 ng/ml) and emricasan (1 μM). Cells were lysed in 1% Triton X-100 lysis buffer (30 mM Tris-HCL pH 7.4, 120 mM NaCl, 2 mM EDTA, 2 mM KCl, 1% Triton X-100 supplemented with protease inhibitors and 10 mM PR619) on ice. Cell lysates were rotated at 4°C for 20 mins then centrifuged at 4°C at 14,000 rpm for 15 mins. 20 μL of protein G Sepharose (SIGMA), previously blocked for 1 hr with lysis buffer containing 1% BSA, were bound with FADD antibody (1.5 mg antibody/mg protein lysate) and were rotated with cleared protein lysates 4 hr at 4°C. 3x washes in lysis buffer were performed, and immunocomplexes eluted by boiling in 60 μL 1x SDS Laemmli buffer.

### Tube assay

Cells were lysed in DISC lysis buffer (20 mM Tris-HCL pH7.5, 150 mM NaCl, 2 mM EDTA, 1% Triton X-100, 10% glycerol) supplemented with protease inhibitors, 1 mM DTT, PR619 (10 mM) and GST-TUBE (50 mg/ml; 50 mg TUBE/mg protein lysate). Cell lysates were rotated at 4 °C for 20 min then clarified at 4°C at 14,000 rpm for 10 min. 25 μL GST beads were added and pull-downs were performed overnight. Beads were washed 3x in wash buffer (50 mM Tris pH 7.5, 150 mM NaCl, 0.1% Triton X-100, and 5% glycerol), and pulled-down proteins eluted by boiling in 50 μl 1x SDS Laemmli buffer.

### Immunostaining

Freshly isolated organs were fixed with 4% PFA overnight, washed with PBS for 24 h, and embedded in paraffin (lung, spleen and skin) or cryopreserved (skin). Paraffin blocks and cryopreserved tissues were sectioned into 3μm thick consecutive thick slices. After standard rehydration, a short antigen-retrieval with 1x sodium citrate pH 6.0 (Sigma-Aldrich C9999) was performed in a microwave for 5 min at 80% power. Tissue sections were then permeabilized with 0.2% (v/v) Triton X-100 in Animal Free Blocker and Diluent (Vector Lab SPS035) at RT for 10 min. Next, the sections were stained for the following antibodies (diluted in 0.2% (v/v) Triton X-100 in Animal Free Blocker and Diluent) overnight at 4°C: CC10 (1:50, Santa Cruz-sc365992), α-smooth muscle actin-Cy3 conjugated (1:50, C6198, Sigma-Aldrich), and CD31 (1:50, 550274 BD Pharmingen). Note that in CD31-stained cryosections antigen-retreival was not performed. Primary antibodies were then visualized by secondary antibodies conjugated to Alexa Fluor 488 (1:200 Thermo Fisher Scientific A-11008) diluted in 0.2% (v/v) Triton X-100 in Animal Free Blocker and Diluent containing 1:1000 Hoechst 33342 for nuclei staining at RT for 1h. For the detection of late PCD+ cells, the ApopTag^®^ Red In Situ Apoptosis Detection Kit (Merck S7165) was used. Briefly, sample sections were washed twice with PBS and treated with equilibration buffer for 10 sec, incubated with the working strength TdT enzyme for 1 h at 37°C, followed by 10 min of stop buffer, and subsequent 30 min of Rhodamine antibody solution at RT. Finally, slides were quickly washed with PBS and mounted with ProLong^™^ Gold Antifade Mountant (Thermo Fisher Scientific P36934). Finally, images were acquired with a confocal fluorescence microscope (Stellaris 5 LIAchroic inverse).

### Intestine histological score

Formalin-fixed and paraffin-embedded intestinal Swiss-rolls were sectioned (3μm) and stained with H&E. Histological evaluation was performed using the scoring system as described in(*60*). In brief, histopathology scores are composed of four parameters: epithelial hyperplasia, epithelial injury, tissue inflammation and epithelial cell death. Histological sub-scores for each parameter: 0, absent; 1, mild; 2, moderate; 3, severe. An “area factor” for the fraction of affected tissue was assigned and multiplied with the respective parameter score (1=0-25%; 2=25-50%; 3=50-75%; 4=75-100%). Each area was scored individually and multiplied with the correlating area factor. Total histology score was calculated as a sum of all parameter scores multiplied with their area factors. Maximum score was 48. Evaluation was performed in a blinded fashion.

### Cell death analysis

Cells were seeded at 8.000 cells (MDFs and LFs), 10.000 cells (LECs) and 50.000 cells (BMDMs) per well of a 96 well plate the day before the experiment. The following day they were treated as indicated in the Fig. legends in the presence of 5 μM of Sytox Green. Live uptake of Sytox Green by dead cells was monitored every hour over a period of 24 or 48 hours via an IncuCyte S3. The percentage of dead cells was calculated by using the Basic Analyzer of the IncuCyte 2020B software and the metric Sytox Object count per well normalized to the area confluence. The positive control (cells treated with TSE) was taken as 100% death and with this the percentage of cell death over time was calculated.

### Protein expression and purification

The pGEX-GST-TUBE construct was transformed into BL21 (DE3) cells and cultured in LB medium at 37°C. Protein expression was induced overnight at 18°C with 0.1 mM IPTG when OD600 reached 0.6. Cells were centrifuged at 4500 rpm for 20 minutes, resuspended in lysis buffer (PBS+300mM NaCl, 1mM DTT supplemented with protease inhibitors) and sonicated 4 times for 30 seconds at maximum amplitude. Lysates were then spun 30min, 4°C, 4500 rpm and cleared lysates were added to GST beads O.N. in rotation at 4°C. Beads were then washed and GST-TUBE eluted using elution buffer (50 mM Tris-HCl pH 8.5, 150 mM NaCl, 1mM DTT and 6mg/ml glutathione) two consecutive times. The elution product was dialyzed using Slide-A-Lyzer cassette in TBS buffer (50 mM Tris pH 7.5 et 150 mM NaC)/1 mM DTT and stored at −80°C.

### Gel filtration

Cellular lysates were separated on a Superose 6 HR 10/30 size-exclusion column and an AKTA Purifier protein purification system (GE Healthcare), essentially as described previously(*61*). Aliquots from each fraction were retained for western blotting, and fractions 12-16, 17–21, 22-26, 27-31, 32–36, 37-41, 42-46 and 47-51 were pooled and used for immunoprecipitation experiments.

### Statistical analysis

The number of independent experiments for each dataset is stipulated in the respective figure legends. Comparisons were performed with a Student’s t test or log-rank Mantel-Cox test (Fig. 7A), whose values are represented in the figures as*P ≤ 0.05, **P ≤ 0.01 and ***P ≤ 0.001using Prism v.8.2 (GraphPad).

## Acknowledgments

We would like to thank all members of the Liccardi, Pasparakis, Kashkar and Peltzer labs for sharing reagents and for the helpful discussions. We also thank the CECAD imaging facility for the support with image acquisition.

## Author contributions

A.A. supervised the study. K.M.L., D.P.S., M.Z., C.C.F. and A.A. designed, performed and analyzed most of the experiments. I.L., K.N., E.J. and N.I. helped performing some of the experiments in Fig. 3 and 4. P.M. provided the *Cflip*^*D377A*^ mouse. I.U., D.d.M., M.A., J.P. M.C.A. and H.W. designed, performed and helped with the analysis of the SARS-CoV infection model. A.C., E.S. and M.M. designed, performed and analyzed the gel filtration experiments. M.P. and S.E. performed the wound healing experiment and helped with the analysis of the results. M.B., X.S., W.T., and A.L. helped with infections models. M.I. and X.H. performed the FACS-based immune-analysis. N.P. contributed intellectually and provided guidance with the *Sharpin^cpdm^* model. The Figures were prepared by A.A., K.M.L., D.P.S. and M.Z. All authors assisted with data interpretation and manuscript editing.

## Supplementary figure legends

**Supplementary Fig. 1.** (**A**) *Cflip*^*f/f*^ and *Cflip^-/-^* and (**B**) *Fadd^WT^* and *Fadd^-/-^* were treated with TNF (100 ng/ml), (**C**) *Caspase3^f/f^Caspase7^f/f^* and *Caspase-3^-/-^Caspase7^-/-^* MDFs were treated with TNF (100 ng/ml) and cycloheximide (1 μg/ml), (**D**) *Caspase-8^WT^* and *Caspase-8^-/-^* MEFs were treated with TNF (100 ng/ml) and cell death was measured over time by calculating the percentage of Sytox Green positive cells. (**E**) Lysates of *Cflip^-/-^* MDFs, reconstituted as shown in the Fig., were immunoblotted with the indicated specific antibodies (EV: empty vector). (**F**) *Cflip^-/-^* MDFs, reconstituted as in E were treated with TNF (100 ng/ml) and cell death was measured over time by calculating the percentage of Sytox Green positive cells.

**Supplementary Fig. 2.** (**A**) Cartoon depicting the *Cflar* exons (upper part). DNA sequence alignment of the region bearing the D>A mutation (lower part). (**B**) Observed and expected numbers of mice of the indicated genotypes from *Cflip^WT/D377A^* intercrosses. (**C**) Weight of 12 weeks old *Cflip^WT^* and *Cflip^D377A^* homozygous mice. (**D**) Aging curves of *Cflip^WT^* and *Cflip*^*D377A*^ homozygous mice. (**E**) FACS analysis of hematopoietic cells isolated from spleen and bone marrow of *Cflip^WT^* and *Cflip^D377A^* homozygous mice (n=4 mice per genotype).

**Supplementary Fig. 3.** (**A**) *Cflip^WT^* and *Cflip^D377A^* MDFs were treated with TNF (100 ng/ml) and cell death was measured over time by calculating the percentage of Sytox Green positive cells. (**B**) MDFs cFLIP WT and D377A stably expressing a doxycycline-inducible HA-tagged CrmA construct and WT MDFs expressing an empty vector (EV) were treated for 72 hours with doxycycline (1μg/ml). Cell lysates were then analyzed by immunoblotting using an anti-HA specific antibody. (**C**) LFs were pre-treated with Enbrel (50 μg/ml) and IFNγ before being subjected to emricasan (1 μM) treatment. Cell death was measured as in A. MDFs WT and D377A mutant were treated with TNF (10 ng/ml) and cycloheximide (1 μg/ml) (**D**), gemcitabine (10 nM) (**E**), paclitaxel (500 nM) (**F**), doxorubicin (1 μg/ml) (**G**) and cisplatin (1 μg/ml) (H), and cell death was measure as in A.

**Supplementary Fig. 4.** (**A**) WT and D377A-mutant MDFs were treated with IFNg (100 ng/ml) and emricasan (1 μM) for the indicated time points and protein extracts were analyzed by immunoblotting with the indicated specific antibodies (n=2). (**B**) WT and D377A mutant MDFs stably expressing a doxycycline (dox)-inducible construct encoding for CrmA were treated for 72 hours with dox and then with TNF (T) (100 ng/ml) for the indicated time points. protein extracts were analyzed by immunoblotting with the indicated specific antibodies (n=3). (**C**) *Cflip^WT^* and *Cflip*^*D377A*^ MDFs were treated with TNF (1 ng/ml) and emricasan (1 μg/ml) for the indicated time points and protein lysates were subjected to TUBE pull-down. The pull-down fractions were left untreated or treated with the deubiquitinase USP2 and analysed by immunoblotting (n=3). (**D**) MDFs and (**E**) BMDMs were treated with TNF (100 ng/ml) for the indicated time points and protein extracts were analysed by immunoblotting with the indicated specific antibodies (n=2).

**Supplementary Fig. 5.** (**A**) WT MDFs were treated for 4 hours with TNF (10 ng/ml), Smac mimetic (250 nM) and emricasan (1 μg/ml) and lysates separated on a Superose 6 column. Aliquots from each fraction were retained and analyzed by immunoblotting with the indicated specific antibodies (1^st^ step) or pooled as indicated and subjected to FADD immunoprecipitation (2^nd^ step). Immunocomplexes were then analyzed by immunoblotting as indicated.

**Supplementary Fig. 6.** (**A**) Spleen of 3 and 7 weeks old mice of the indicated genotypes (left), with the respective weight express as percentage of total body weight (right). (**B**) K6 (green) staining on skin section of 3 months old *Sharpin^WT^* and *Sharpin^cdpm^* mice. Scale bar 100 μm. (**C**) Small intestine sections of *Sharpin^WT^* and *Sharpin^cdpm^* mice. Arrowhead indicates a Peyer’s patch. Scale bar 1 mm.

