## Supplementary figures for "Cleavage of cFLIP restrains cell death during viral infection and tissue injury and favors tissue repair"

Supplementary figure 1

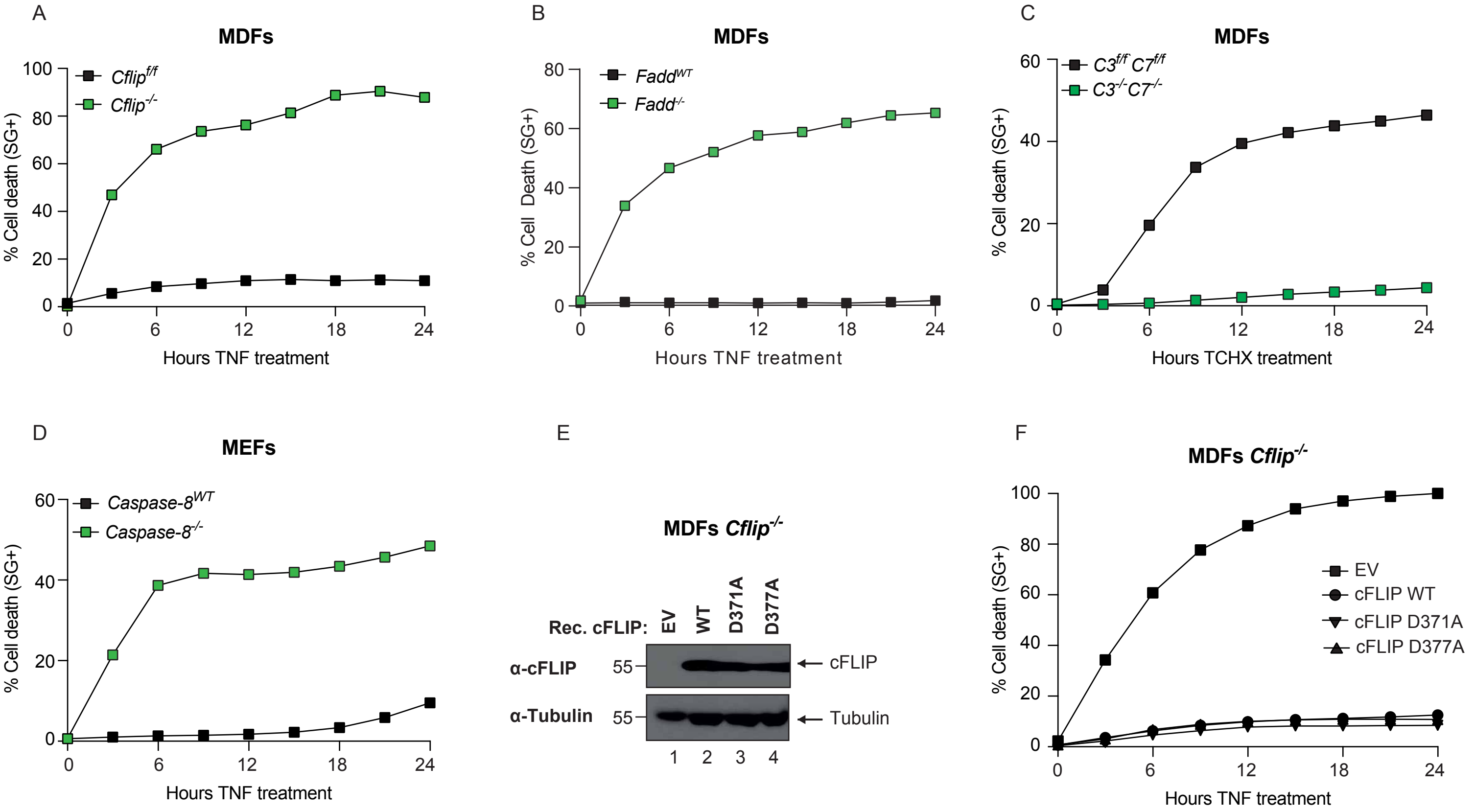

### Supplementary figure 2

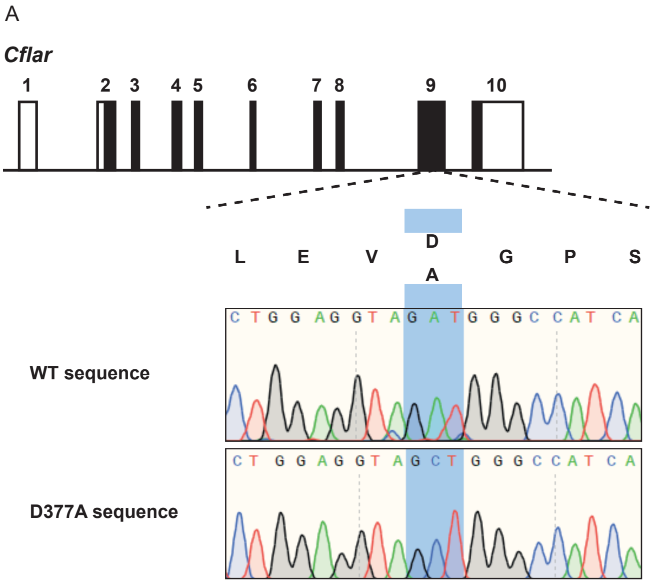


B

| Genotype | # of mice | Observed (%) | Predicted (%) |
|---|---|---|---|
| *Cflip*$^{WT/WT}$ | 60 | 26,5 | 25 |
| *Cflip*$^{WT/D377A}$ | 112 | 49,5 | 50 |
| *Cflip*$^{D377A/D377A}$ | 54 | 24 | 25 |

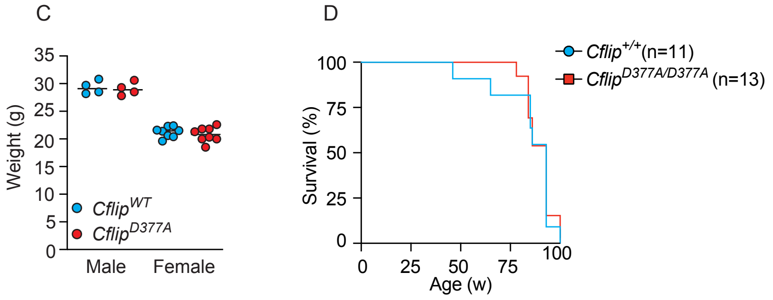


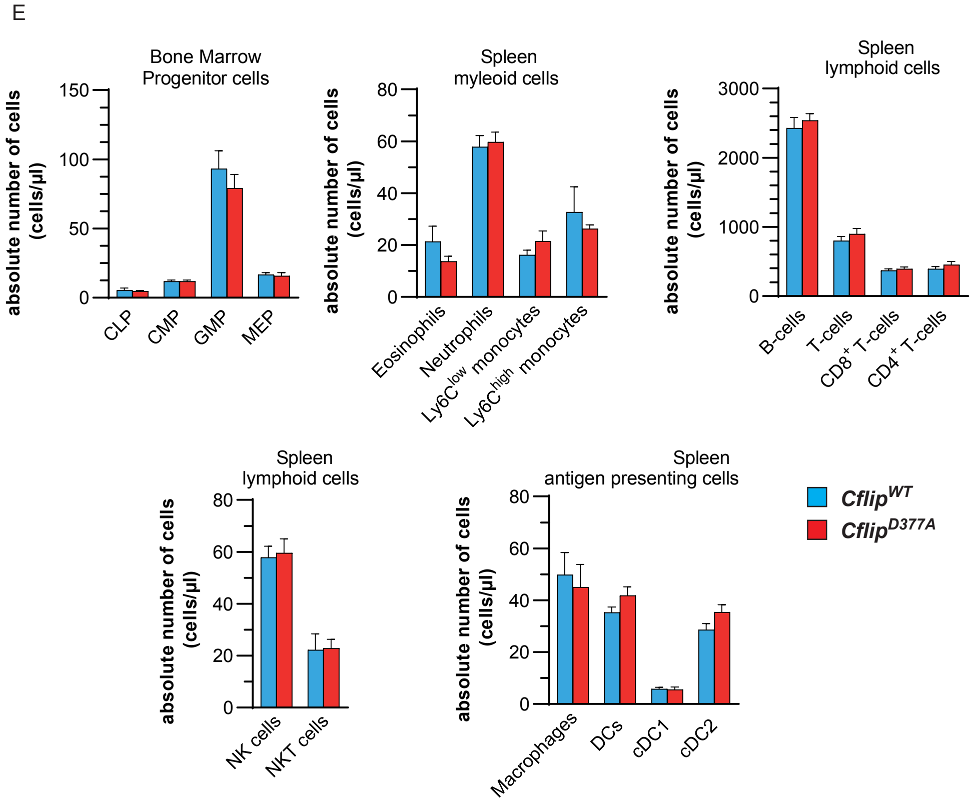

Supplementary figure 3

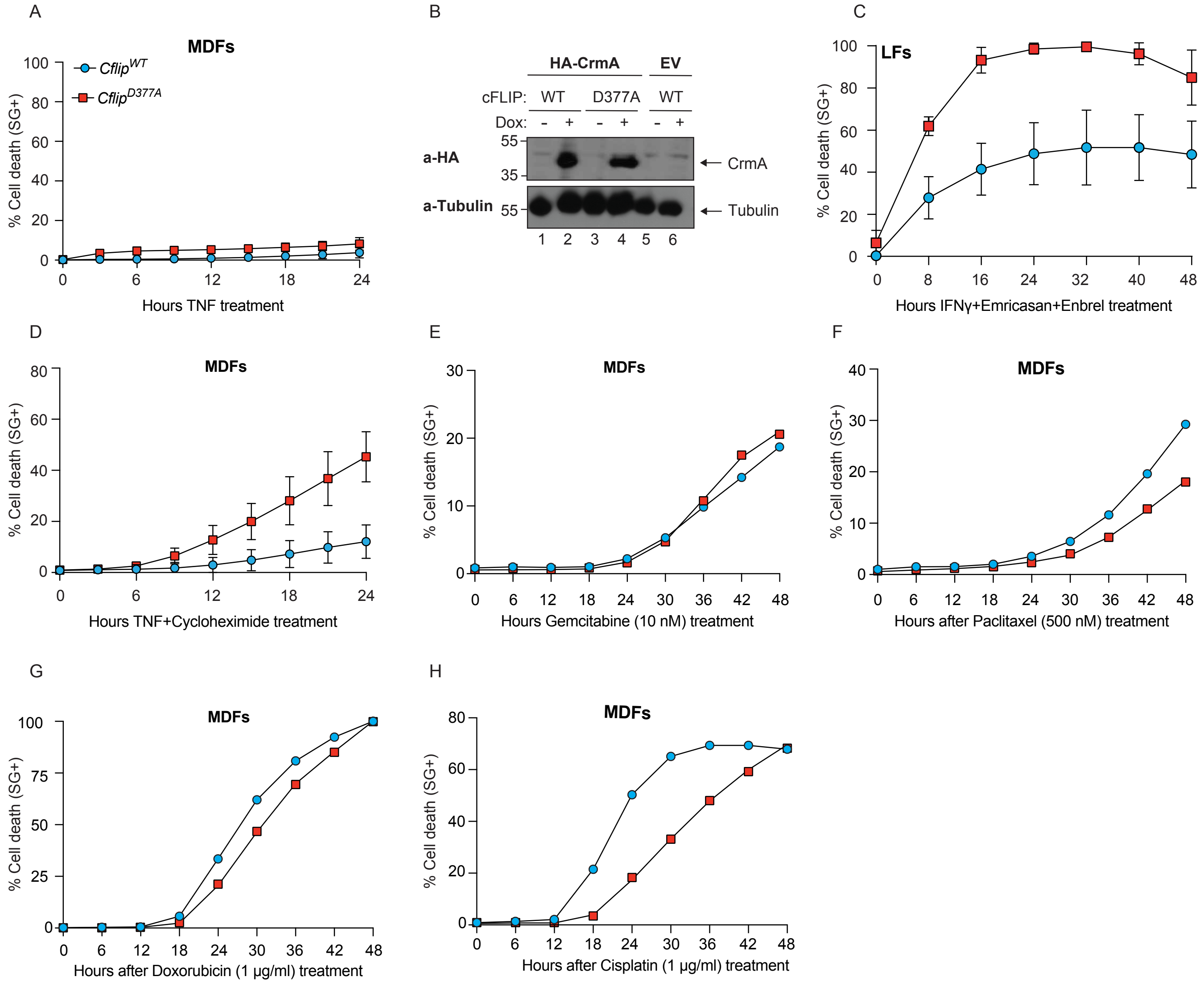

Supplementary figure 4

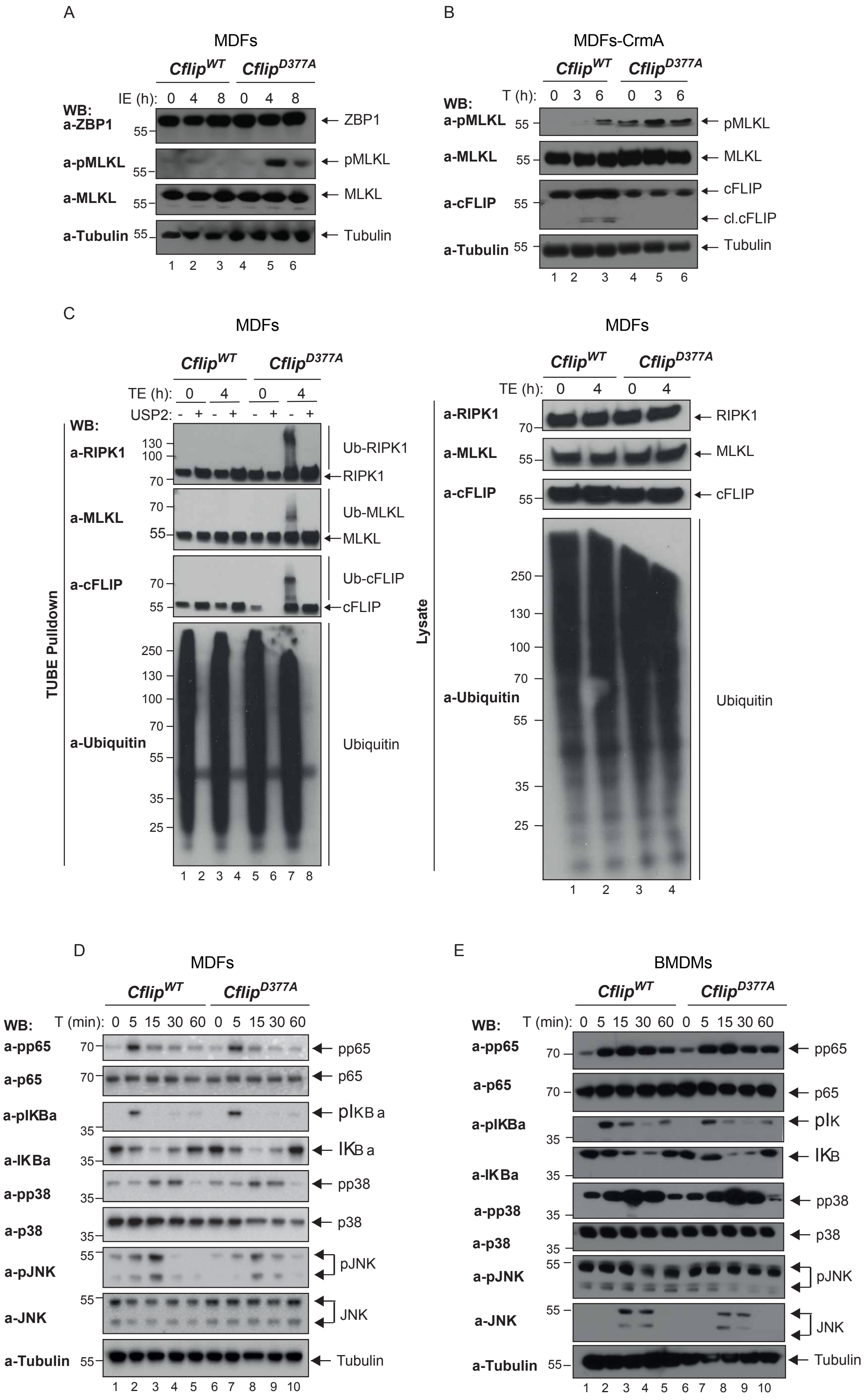

### Supplementary figure 5

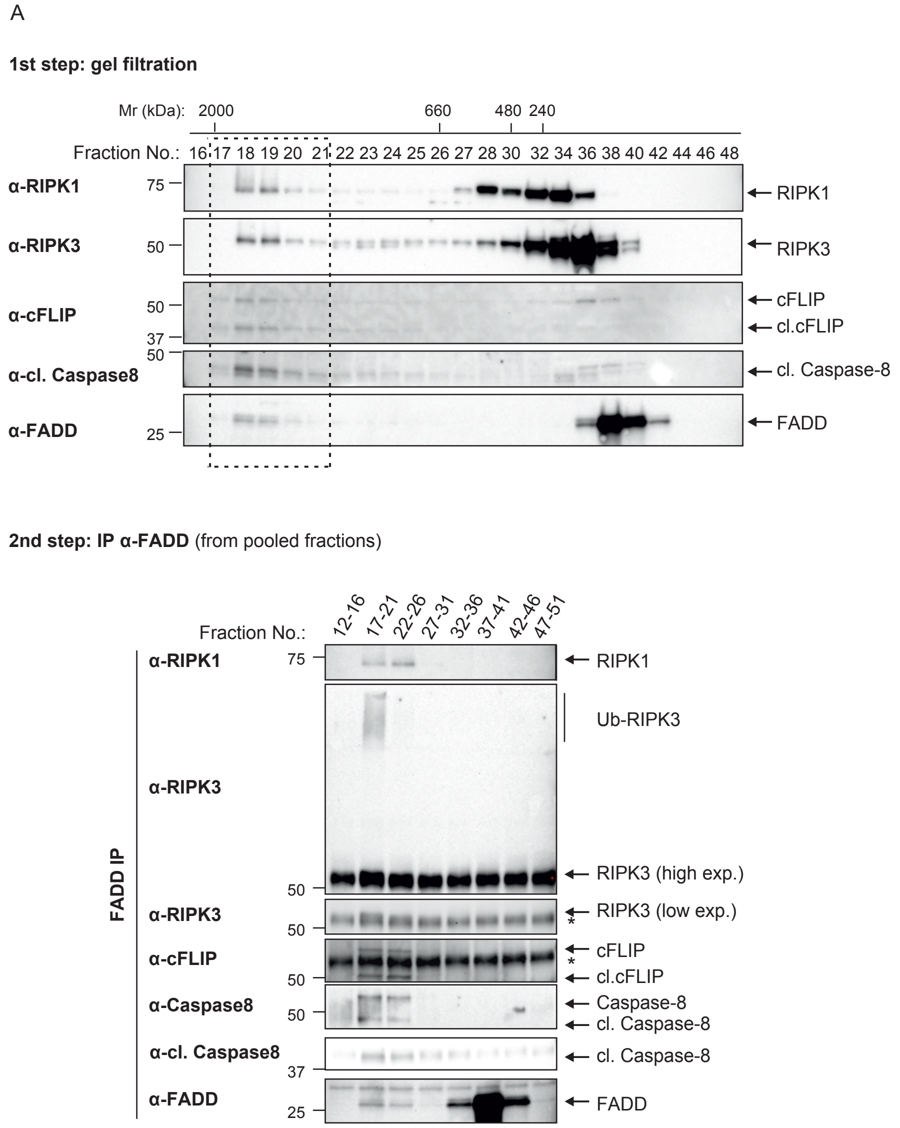

### Supplementary figure 6

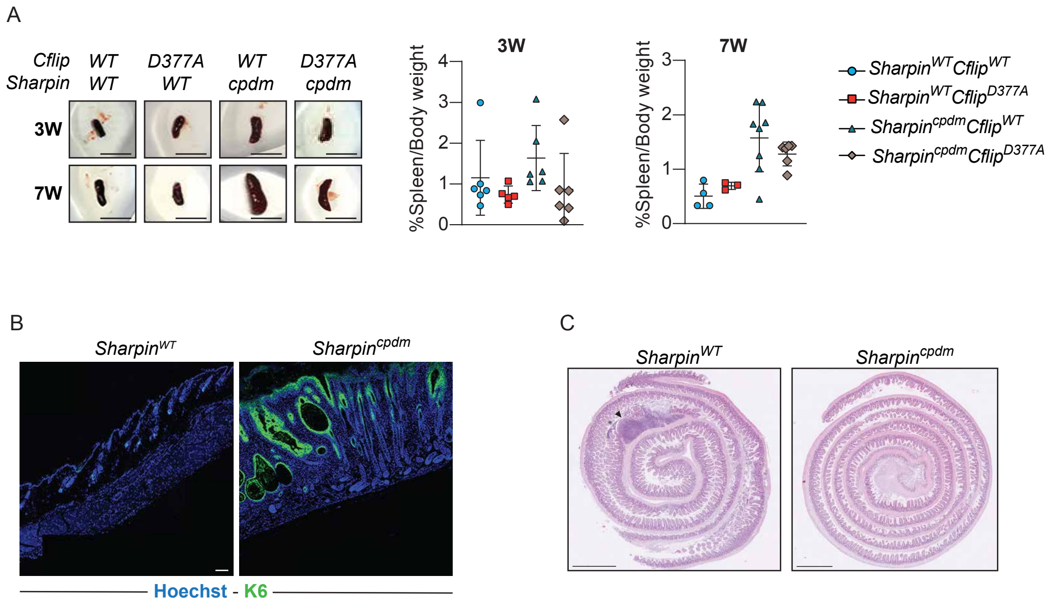
